# CPSign - Conformal Prediction for Cheminformatics Modeling

**DOI:** 10.1101/2023.11.21.568108

**Authors:** Staffan Arvidsson McShane, Ulf Norinder, Jonathan Alvarsson, Ernst Ahlberg, Lars Carlsson, Ola Spjuth

**Affiliations:** Department of Pharmaceutical Biosciences and Science for Life Laboratory, Uppsala University, Sweden; Department of Computer and Systems Sciences, Stockholm University, Sweden; MTM Research Centre, School of Science and Technology, Örebro University, Sweden; Department of Computing, Jönköping University, Sweden; Department of Computer Science, Royal Holloway University of London, UK

## Abstract

Conformal prediction has seen many applications in pharmaceutical science, being able to calibrate outputs of machine learning models and producing valid prediction intervals. We here present the open source software CPSign that is a complete implementation of conformal prediction for cheminformatics modeling. CPSign implements inductive and transductive conformal prediction for classification and regression, and probabilistic prediction with the Venn-ABERS methodology. The main chemical representation is signatures but other types of descriptors are also supported. The main modeling methodology is support vector machines (SVMs), but additional modeling methods are supported via an extension mechanism, e.g. DeepLearning4j models. We also describe features for visualizing results from conformal models including calibration and efficiency plots, as well as features to publish predictive models as REST services. We compare CPSign against other common cheminformatics modeling approaches including random forest, and a directed message-passing neural network. The results show that CPSign produces robust predictive performance with comparative predictive efficiency, with superior runtime and lower hardware requirements compared to neural network based models. CPSign has been used in several studies and is in production-use in multiple organizations. The ability to work directly with chemical input files, perform descriptor calculation and modeling with SVM in the conformal prediction framework, with a single software package having a low footprint and fast execution time makes CPSign a convenient and yet flexible package for training, deploying, and predicting on chemical data.

## Introduction

Ligand-based modeling and quantitative structure-activity relationships (QSAR) are computational methods used in drug discovery to predict properties of small molecules, such as binding affinity or activity towards a protein target, and toxicity [1, 2, 3]. The approach relies on the structure and properties of known chemical structures, and commonly takes advantage of machine learning to construct predictive models. Over the years the available data in public repositories related to cheminformatics have increased, and the applications and accuracy of predictive models have expanded and improved. This has lead to an increased utilization of ligand-based modeling in drug discovery projects [4].

The predictive performance of machine learning models is commonly measured on an external test set or using cross-validation, with accuracy, AUC, F1 scores (classification), and RMSE and R^2^ (regression) as commonly used metrics. However this does not relay the level of confidence for individual objects predicted. When predicting two different objects, it would seem natural that the object that is most dissimilar compared to the training data would result in a larger prediction interval to reflect greater uncertainty, and vice versa. In drug discovery, where predicted objects in many cases are novel chemical structures, this is particularly important, and concepts and approaches to determine a model’s applicability domain have been proposed to this end [5]. However in most cases these are ad hoc methods without a proven theoretical underpinning.

Conformal prediction is a framework that provides a way to generate valid prediction intervals for a wide range of machine learning algorithms [6]. Unlike traditional prediction intervals, which applies the same certainty regardless on the predicted object, conformal prediction constructs prediction intervals that are both guaranteed to be valid and based on the estimated difficulty of the predicted objects. This makes conformal prediction a powerful tool for machine learning in settings where the underlying distribution of data is unknown, and a way to address the applicability domain assessment for compounds [7, 8].

Conformal prediction has been extensively used in drug discovery [9] with applications including screening [10], toxicology prediction [11, 12], property prediction [13], target prediction [14], and prediction of pharmacokinetics [15, 16]. More recently, conformal prediction has also been used with Deep Neural Networks in drug discovery applications [17, 18, 19] and in medical applications [20].

Existing software for conformal prediction include the Nonconformist software [21] which is a Python implementation of the conformal prediction framework that has been used in many drug discovery projects [22, 23, 24]. Crepes [25] is a more recent Python package for generating conformal regressors and predictive systems. For a more extensive list of resources, papers, and software related to conformal prediction, we refer to the Awesome Conformal Prediction GitHub repository [26].

In this manuscript we present CPSign, a standalone software tool that implements conformal prediction for cheminformatics modeling. We start by introducing conformal prediction and Venn-ABERS prediction as well as the default CPSign methods; Signatures [27, 28] for molecular representation, and support vector machines (SVMs) for machine learning modeling. We continue to discuss the implementations in CPSign and associated tools, and also present a comparison between CPSign and several other methods for a set of regression and classification datasets.

## Methods

### Conformal prediction

Conformal prediction is a mathematical framework built up by a collection of algorithms used for producing confidence guarantees for standard machine learning algorithms [6]. There are many resources that has introduced it in different settings, e.g., in the drug discovery domain [9, 29]. Here we make a brief introduction to the subject, focusing mainly on the *inductive* versions of the algorithms, in which an underlying scoring algorithm is trained once and then later reused for all future predictions (until enough new training data has been accumulated to warrant a full retraining to include new knowledge in the model).

At the heart of conformal prediction lies the notion of *nonconformity*, which intuitively is a measure of how “strange” an object is compared to other objects. The term *object* here refers to the features *x* of a training observation (*x*,*y*), where *y* is the label of the observation. The nonconformity of an object *i* is often referred to as *αi* and computed using a *nonconformity function*; *h*(*xi*). This function, *h*, is based on the output of an *underlying scoring algorithm* - which could be any machine learning algorithm that produces a prediction score. In the inductive setting, where the underlying algorithm is trained only once, the available training data is split up into two disjoint sets; the *proper training* and the *calibration* sets. The proper training set is used for training the underlying algorithm, while the calibration set is used for calibrating the predictions - based on the nonconformity function.

The classification and regression algorithms differ slightly and we refer readers to e.g. Vovk et al. [6] for a detailed explanation of them. But in essence the classification algorithm computes the nonconformity, α*_i_*, for all *n* instances in the calibration set during training, resulting in a list *α*_1_,…, *α_n_*. When predicting a new test-object, *x_n_*_+1_, the nonconformity, α*_n_*_+i_, is calculated and then ranked against the list *α*_1_,…, *α*_*n*+1_ (i.e., including the *n* + 1 test instance) according to Equation 1, resulting in a p-value for the object. This ranking is in most cases performed separately (referred to as *mondrian*) for each possible class label *y* (i.e., with a separate list of α values for each class). The resulting prediction of a test-object is a set of p-values, one for each possible class label. These p-values can then be subjected to a statistical test in order to obtain a set-prediction, i.e. if we wish to have 90 % confidence in the prediction we specify a significance level, ε, of 0.1 and have to include all labels with a p-value equal to or higher than 0.1 in the prediction set. The resulting prediction sets can thus be empty (no classes predicted), single-label (informative) or multi-label (less informative).

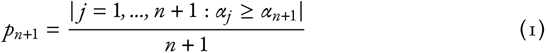

In the regression algorithm it is common to use a nonconformity function that also attempts to scale the prediction intervals based on the predicted difficulty of the test-object, commonly performed by training an additional *error model* that is trained on e.g. the residuals produced when predicting the training set. The regression algorithm, in contrast to classification, also require the user to specify a desired significance level (ε) at prediction time and the output is a prediction interval for the given *ε*. This prediction interval should enclose the true label, *y*, with a probability of 1-ε or greater (i.e. the expected error is at most ε). Naturally, as the desired significance level is decreased the predictor has to increase the prediction interval in order to comply with the lowered level of accepted errors (see Figure 1 for an illustrative example).

**Figure 1:**
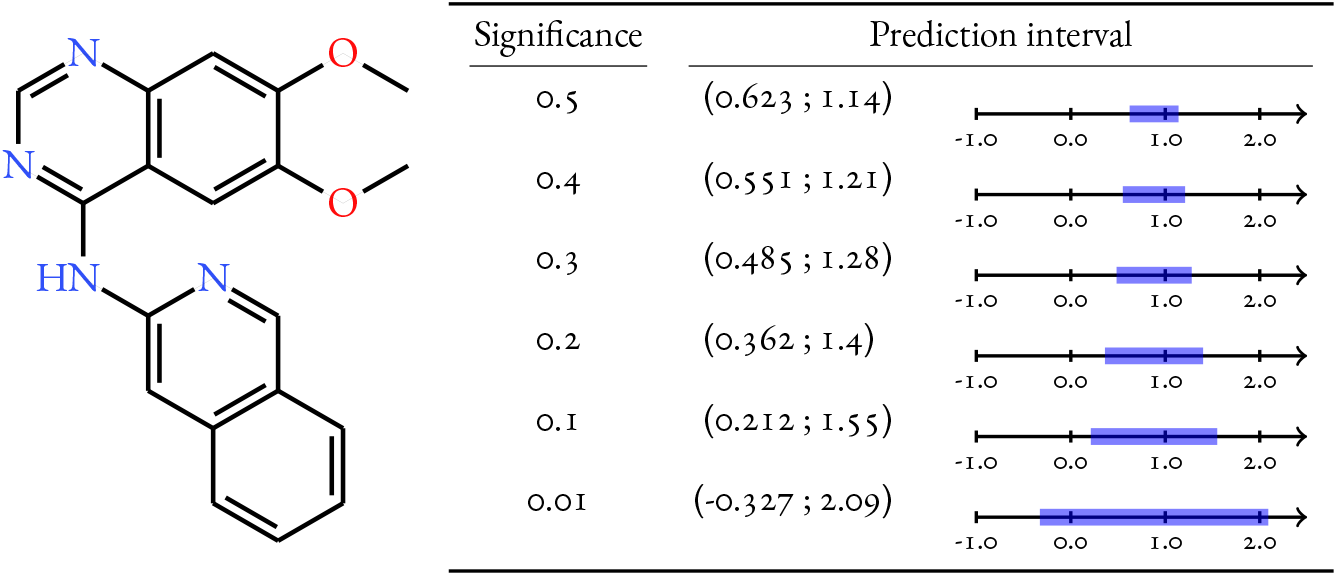
Predictions made with the lipophilicity model from the evaluation using different significance levels. Selecting a lower significance, to be more certain that the true prediction is included in the predicted interval, leads to a larger interval.

Under the relatively week assumption of *exchangeability* of calibration and test-data, these inductive versions of conformal predictors are proven to produce *valid* (well-calibrated) predictions, i.e., that the error rate is equal to or smaller than the specified significance level [6]. Furthermore, in the case of classification, given that a mondrian (class conditional) calibration is used, the guarantee holds individually for each class and has been shown to handle imbalanced datasets very well without the need to apply balancing techniques [30, 31, 18]. However, this guarantee may in practice be difficult to achieve sometimes due to assay drifts [12] or in case data splitting is performed in a non-random way (such as scaffold splitting). The validity is thus commonly assessed by calculating the error rate for a set of significance levels, or by plotting a *calibration curve* of error rate vs significance level across a range of significance levels (see e.g. Figure 3A).

**Figure 2:**
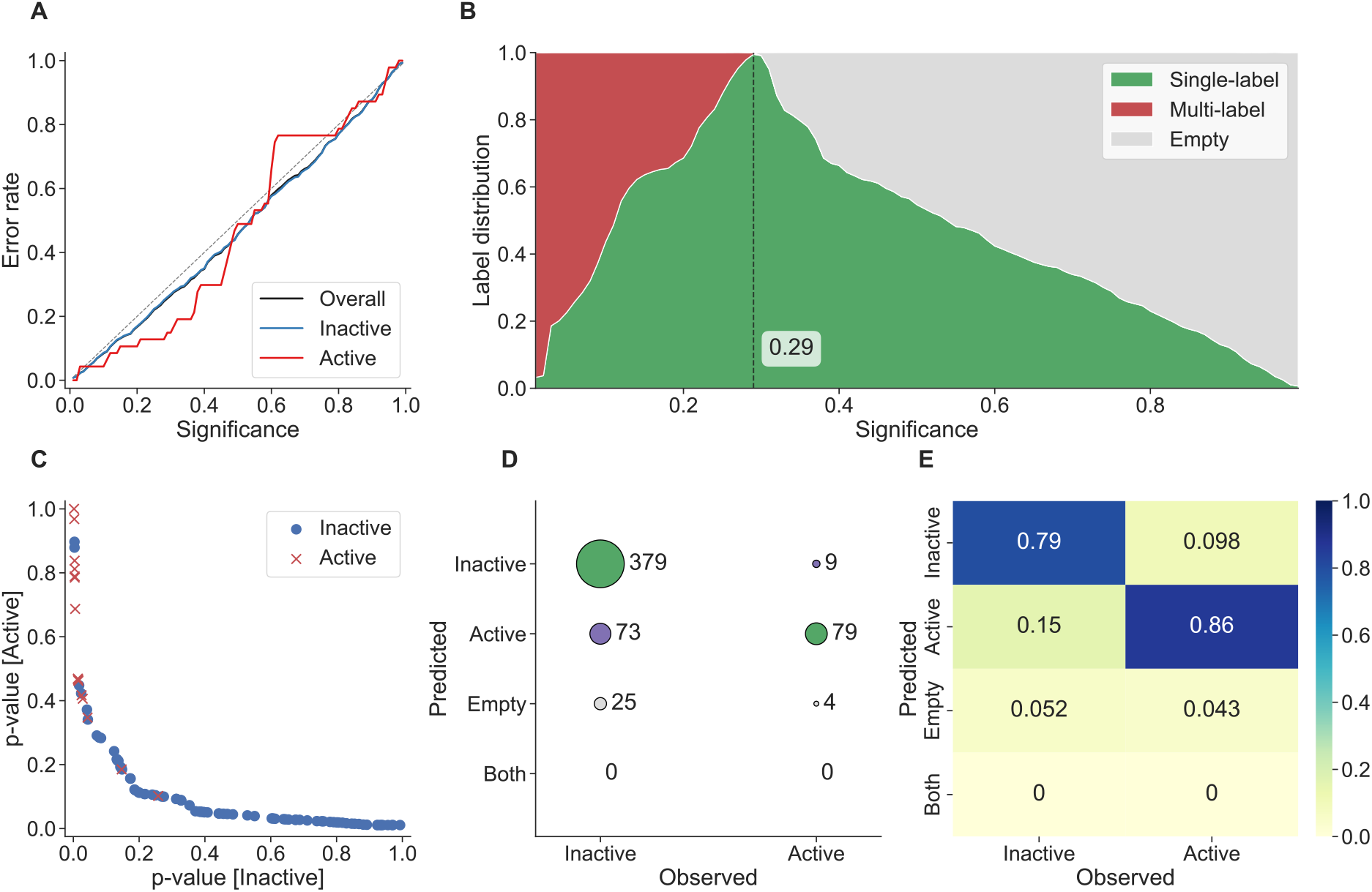
Figures showing features from conformal classifiers. Panel A displays the calibration curve from the difficult NR-AR-LBD dataset used in the evaluation, having only 3.5 % active instances and proved problematic for all tested methods in predicting the minority class. For a well calibrated model, both classes’ error rates should be less than or equal to the significance level and thus follow the gray dashed diagonal line in the chart. The error rate of the inactive class follows the diagonal well, likewise the black (mostly covered behind the blue line) of the overall error rate. This shows one of the benefits of conformal prediction, the possibility to inspect calibration independently for each class. Panel B shows the distribution of prediction sets across all significance levels, which can be used for finding the best significance level to use for a specific data set and to analyze the predictive efficiency of the model. Panel C displays a plot of the two p-values against each other for 100 randomly picked predictions for the SR-MMP dataset, showing the possibility to easily find potential outliers such as the two inactive (blue circles) with high p-values for the active class and low p-values for the inactive class. This plot can also be used to find test objects that are predicted with low p-values for both classes (lower left area), constituting a region with predictions of lower confidence. Panel D and E shows the conformal version of a confusion matrix and normalized heatmap for a fixed significance level of 0.2, where predictions can also be empty sets as well as multi-label predictions. All figures were created using our Plot_utils python library described later.

**Figure 3:**
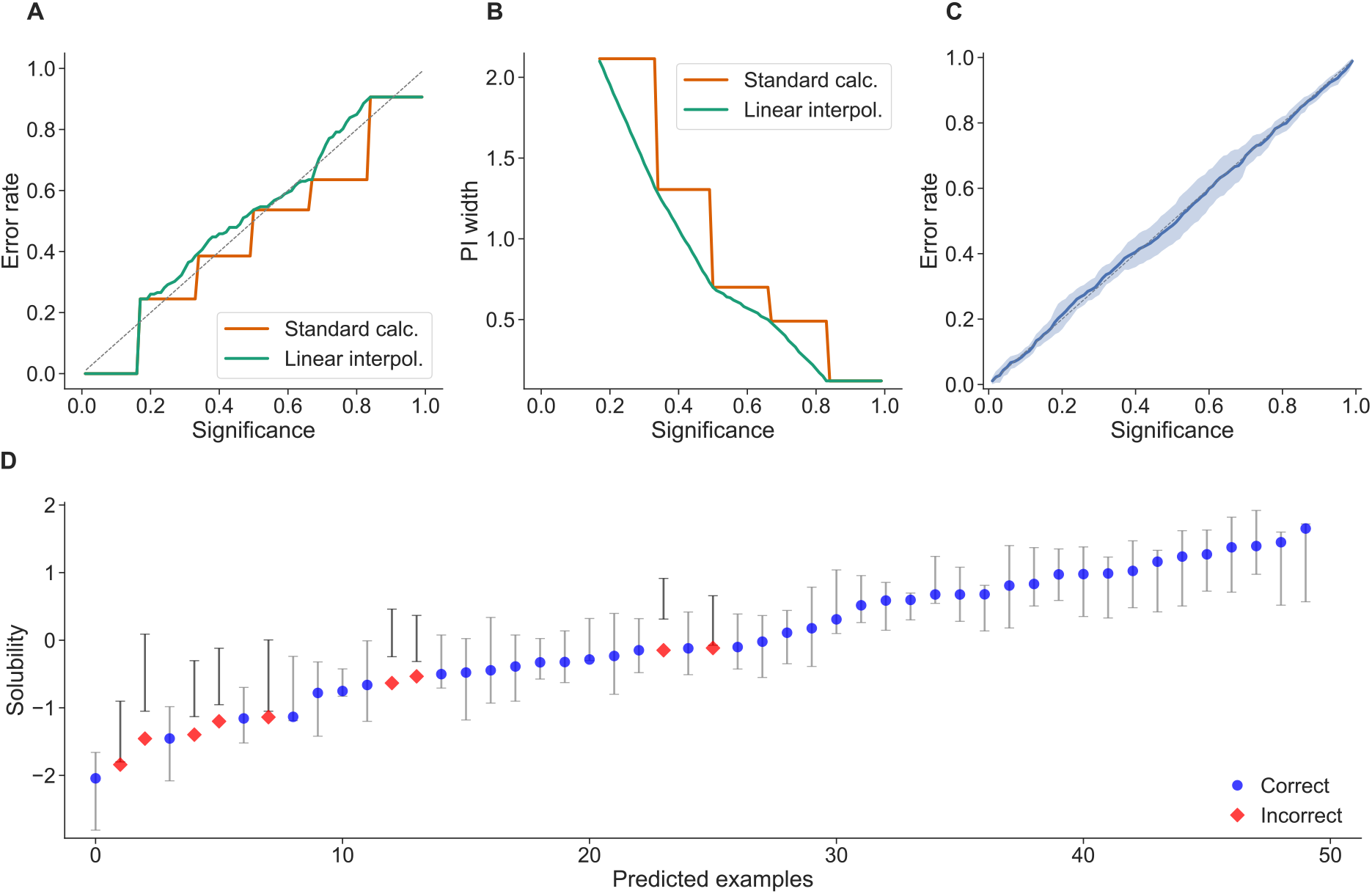
Figures showing features from conformal regression predictors. Panel A and B displays the performance of a conformal regression predictor with only five instances used in the calibration set, without (orange) and with (green) interpolation used in the calculation. Without interpolation (standard calculation) the prediction intervals can only change at five significance levels and give rise to sharp steps, whereas the linear interpolation can reduce the influence of having a small calibration set and give smoother changes in the prediction intervals. Further note that in panel B the curves start at 0.17 as the prediction intervals are (−∞; ∞) before that. Panel C displays a calibration plot of the result of a 10-fold cross-validation on the train set of CHEMBL205_pKi, where the shaded area is +/- the standard deviation from the 10 folds. Panel D displays 50 randomly picked predictions made by the CPSign model trained on the ESOL dataset from the evaluation, showing the possibility to easily analyze individual predictions and e.g. find patterns in erroneous predictions. All figures were created using our Plot_utils python library described later.

Once calibration has been assessed, the goal is to produce as informative predictions as possible, referred to as *predictive efficiency*. A conformal predictor could always place all possible class labels in the prediction set or predict the interval (-∞; ∞), and thus always be correct - but those predictions are not informative. The predictive efficiency thus relates to how specific the predictions are (small prediction sets or tight prediction intervals). Many metrics has been proposed for evaluating predictive efficiency, most which are summarized in Vovk et al. [32]. Some of the most commonly used metrics for classification are: observed fuzziness, the average number of predicted classes (average C), and the ratio of single-label prediction sets; where the two latter ones requires a fixed significance level. For regression the most commonly used metrics are the median or mean prediction interval width at fixed significance levels.

Evaluating a conformal model requires more effort and thought than standard point predictions, as we also need to apply domain knowledge into how each dataset should be evaluated. More specifically, we need to choose a sensible level of confidence in the evaluation - something that require insight in what level of confidence is needed for a prediction to be useful, or contrary, how specific must the predictions be to be useful (e.g. tight prediction intervals). An example of this is choosing to use the proportion of single label predictions produced by a classification model at a fixed significance level, which seems sensible as the most useful prediction is when it is of a single label, but *a priori* it may be hard to decide on what confidence level to use. See Figure 2B, where the green area shows the proportion of single-label predictions at any given significance level. If we in advance chose to use 0.2 or 0.3 as significance level the predictions will include many empty prediction sets - the predictive efficiency is actually better at a lower significance level (0.15 being the best), which you may miss in case focusing on a fixed significance level.

### Venn-ABERS Prediction

Apart from the conformal predictors introduced in the previous section, CPSign also supports probabilistic modeling using the Venn-ABERS predictor (VAP) [33, 34]. This algorithm, contrary to the conformal predictors, output probability estimates rather than p-values, which is preferred by some users. The VAP is a special type of Venn predictor which relies on a machine learning model to produce so-called Venn taxonomies. VAP is a multi-probability predictor for binary classification tasks, giving two probability estimates (*p*_0_ and *p*_1_) for each test-object. One of these estimates is the true probability of the test-object, but we do not know which one. This simplest version of VAP relies on splitting the full training set into a proper training and calibration set (called Inductive Venn-ABERS Predictor, or IVAP), in the same manner as the inductive conformal predictors discussed in the previous section. The proper training set is similarly used for training the underlying scoring algorithm and the calibration set is here used for producing the predictions using an isotonic regression where the test-object is included. The calibration step is performed two times, once for each of the possible class labels, where the test-object is augmented with one of the class labels as a tentative label.

An extension to the one-split VAP is to train a Cross-Venn-ABERS (CVAP) [34] model, in which the training set is split several times in a folded fashion similar to *k*-fold cross validation, where an IVAP is trained for each such split. The benefit of this strategy is that the *k* multi-probabilities can be aggregated into a single probability with conditional guarantees [34]. A benefit of VAP is that it has guarantees for producing well-calibrated probabilities, without introducing further assumptions on the data being modeled, something that is not guaranteed by most probabilistic models [35]. In many cases a probabilistic predictor is favorable over a conformal classifier, mostly as probabilities are easier to interpret than p-values, and having the possibility to measure and compare its performance using standard evaluation metrics. However, there are cases where the conformal algorithms are preferable, such as for imbalanced datasets where the mondrian calibration handles the imbalance without requiring balancing techniques - whereas VAP generally needs data balancing or other techniques to perform well. Conformal classification also handle the case of multi-class data, whereas the VAP is only defined for binary classes. For the scenario of dealing with small datasets the transductive conformal predictor may be a more favorable alternative as no valuable training observations must be set aside for the model calibration, whereas the VAP algorithms require a separate calibration set.

In life science research, VAP has been applied in drug screening [36], to predict metabolic transformations [37], and to assess cardio-vascular risk based on in vitro assay data [38].

## CPSign

This section will briefly go through the implementation choices made when developing CPSign and some of the key features, a summary of these can be seen in Table S1.

### Molecular representation - descriptors

Representing chemical structures as numerical features is generally referred to as molecular or chemical descriptors [39], but sometimes also chemical fingerprints although this is mostly used for structural comparisons and similarity searching. Many descriptor implementations have been proposed, with varying performance. The simplest ones include physicochemical descriptors such as molecular weight, number of rotatable bonds, lipophilicity etc. Over the years, a type of topological (2D) fingerprints describing the local environment around each atom, referred to as circular fingerprints [40], have emerged as robust descriptors that sustain efficient modeling with machine learning methods. Several different approaches and implementations exist, with Morgan fingerprints [41], extended-connectivity fingerprints (ECFP) [42], and Signatures [28] being the most widely used. These descriptors can be rapidly calculated and since they stem from chemical substructures, they allow for chemically relevant feature interpretations.

CPSign implements Signatures as the main descriptor type, but also CDK molecular descriptors [43] including ECFPs. The user can also generate descriptors by other tools and load them as properties from CSV or SDF files together with the chemical structures. Additional descriptors can be calculated by extending an interface as explained in section Adding custom extensions.

### Underlying scoring algorithms

CPSign includes the Java versions of the popular LIBLINEAR [44] and LIBSVM [45] packages, both implementing support vector machines (SVMs) allowing for sparse input data. The need for sparse data representation is essentially required when using the Signatures descriptor as the number of features can be in the order of hundreds of thousands for larger datasets - which would require vast amounts of RAM for dense representations. For smaller datasets the RBF-kernel SVM from LIBSVM is preferable (other kernels are also possible to use), but the training time scales poorly for larger datasets [46]. For larger datasets the linear kernel SVM from LIBLINEAR is often preferred, as it also includes heuristics in order to speed up training time [44]. Earlier work has been aimed at finding good sweet spot hyper-parameters for the combination of using the Signatures descriptor with SVMs [47], leading to robust results using the default settings in CPSign.

LIBLINEAR also has a sparse implementation of the logistic regression algorithm that can be used within CPSign, albeit less tested and may require more tuning to achieve good results. Similarly as for the chemical descriptors, users can implement and expose their own machine learning methods to be used as underlying scoring algorithms. We have also made such an extension by wrapping the DeepLearing4j library [48], that can be found in the CPSign-DL4j GitHub repository (https://github.com/arosbio/cpsign-dl4j). Note however that this extension requires the user to build it and adjust it to work for a particular platform and hardware as it requires to run native code.

### Predictor types

CPSign implements the transductive conformal predictor (only for classification), a variety of inductive conformal predictors (both for regression and classification) and the Cross Venn-ABERS predictor (binary probabilistic classifier). For the conformal models there are three base types; TCPClassifier, ACPClassifier and ACPRegressor. The ACPClassifier and ACPRegressor predictors can be changed between running one-split ICP, several splits (ACP [49]) or folded splits (CCP [50]), depending on the data sampling strategy. There is also the option of using a pre-defined split from the user or adding custom splitting strategies such as in Arvidsson McShane et al. [51]. For the TCPClassifier there is no splitting of data into proper training and calibration sets, instead the underlying scoring model is trained once for every possible label *for every test example*. The TCP model can thus use all available data both for training the underlying model and for calibration, but at the cost of being highly computationally demanding, and is thus only recommended for smaller datasets.

For the conformal predictors the notion of nonconformity measure (NCM) is central, and CPSign comes with four NCMs for classification and four for regression (see Table S1). Depending on what the function requires, they can be combined with different types of underlying scoring models, e.g., the InverseProbability requires probability scores from the underlying model and thus restricts the number of available scoring models, and the NegativeDistanceToHyperplane requires the use of SVM as scoring model. For the regression algorithm the NCM also dictates whether an additional error model should be trained in order to predict the difficulty of a test example in order to normalize the prediction intervals. By default the error models will use the same algorithm and hyper-parameter settings as the main scoring model, but it is possible to, e.g., use a more complex (RBF kernel SVM) for predicting the midpoint and then use the computationally cheaper linear kernel based SVM to normalize the prediction intervals with.

Another feature of CPSign is the possibility to use different ways of calculating and handling the p-values, apart from the standard calculation (Equation 1), also allowing to calculate “smoothed p-values” [6], and both linear and splines interpolation [52, 53]. The interpolation options can be useful especially when having small datasets, where only a few examples can be set aside to be used in the calibration set, see Figure 3A-B for a comparison between the standard p-value calculation and linear interpolation when only having five calibration instances. This example is exaggerated and we do not recommend using only five examples for calibration, but shows the usefulness of including interpolation.

### Hyper-parameter tuning

CPSign has robust predictive performance using the default parameters (see the method evaluation section for a comparison against tuning of hyper-parameters, as well as other popular modeling methods). To further improve the model performance it is possible to fine-tune the hyper-parameters using a standard grid search algorithm. For some parameters, e.g., the SVM cost parameter, there are default values to try out - for other hyperparameters the user has to decide which values to evaluate in the grid. There is flexibility in how this should be done, e.g.; choice of performance metric and the evaluation strategy to use (see section Validation strategies). Furthermore, it is possible to choose whether to tune the parameters based on the underlying scoring model by itself, or if the evaluation should be performed based on a conformal or Venn-ABERS predictor. This latter concept can be especially useful if aggregating several ICPs, or if the final model will be a TCP (requiring re-training the underlying scorer model multiple times for each test-prediction), where considerable computational time can be saved.

### Validation strategies

Three validation strategies and several settings thereof are available; *k*-fold cross-validation, single test-train split, and leave-one-out cross-validation. The two former has several configurable parameters such as performing the splits stratified (for classification), the fraction of test-instances, the *k* in *k*-fold, and number of repetitions. Validation strategy is another feature that is extendable with CPSign.

### Interpretation of predictions

CPSign can render images of molecules with an interpretation of the prediction based on the algorithm outlined in Ahlberg et al. [54] (Figure 4). This interpretation is based on feature importance’s that can be mapped back to atoms in the molecule, and can either be in terms of which molecular signature had the highest impact on the prediction (Figure 4A) or as a complete molecule gradient where all features individual contributions are aggregated and mapped back to their originating atoms (Figure 4B). This feature has been especially appreciated by chemists utilizing the predictive models, as it allows for editing chemical structures in a drawing graphical user interface and immediately visualize how the predictions change.

**Figure 4:**
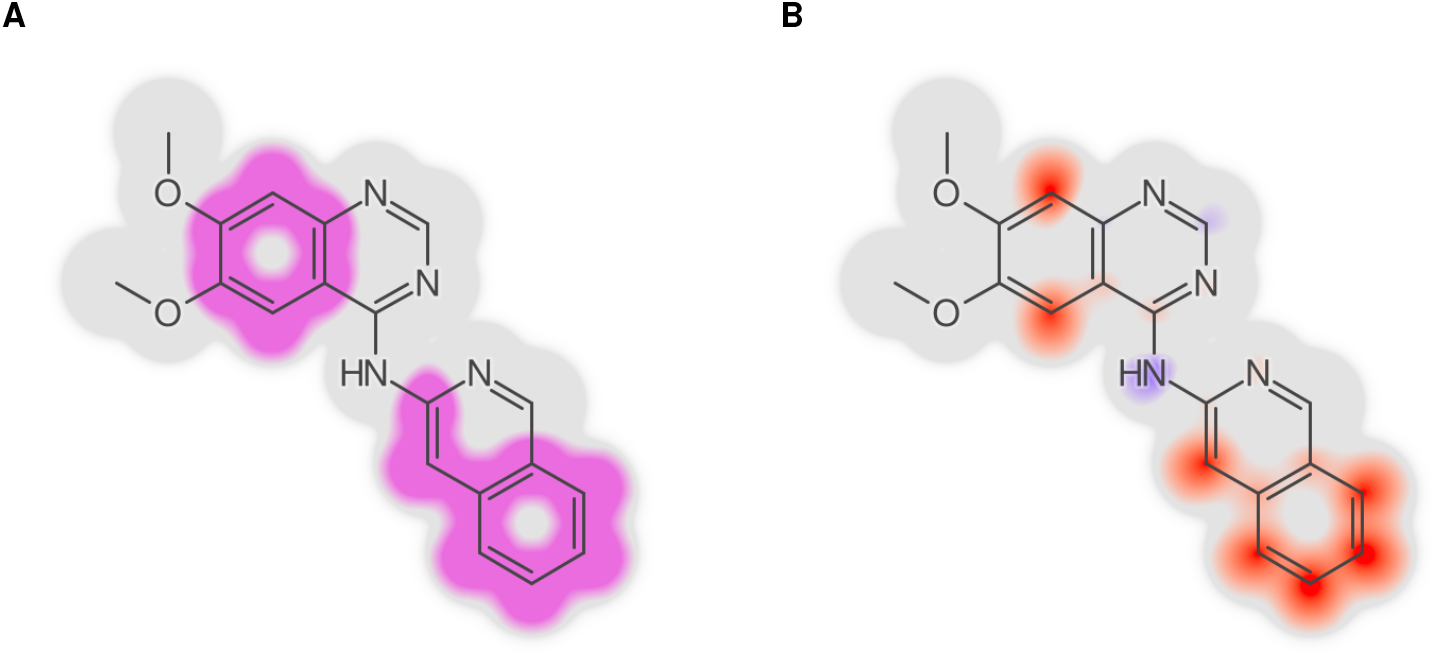
Using the Signatures descriptor allows to map feature importance back to the atoms they originate from, allowing visual interpretations of the predictions, as described in Ahlberg et al. [54]. This can both be visualized as depicting the signature having the largest contribution (the light blue “significant signature”, panel A), or by aggregating the contribution of all signatures to see each atoms’ individual contribution (panel B). These figures are generated using the CPSign method trained on the Lipophilicity data set from the evaluation section and the signature that had the highest contribution was “`[C](p[C]p[C]`)”, which maps to several atoms and thus appears at several locations in the molecule.

### Implementation details

CPSign is written in Java and is thus platform-independent, only requiring a Java runtime of version 11 or later. Build and dependency management is handled by Maven, and published artifacts are available from the Maven central repository for easy inclusion in other JVM-based projects. The code base is split up into several child-modules, outlined in Figure 5B, to allow users to only depend on the parts needed for their particular requirements. For instance, users modeling non-chemical data can do so by depending on the ConfAI module and reduce the dependency graph, and the REST servers depend on the CPSign-API module as no CLI functionality is required.

**Figure 5:**
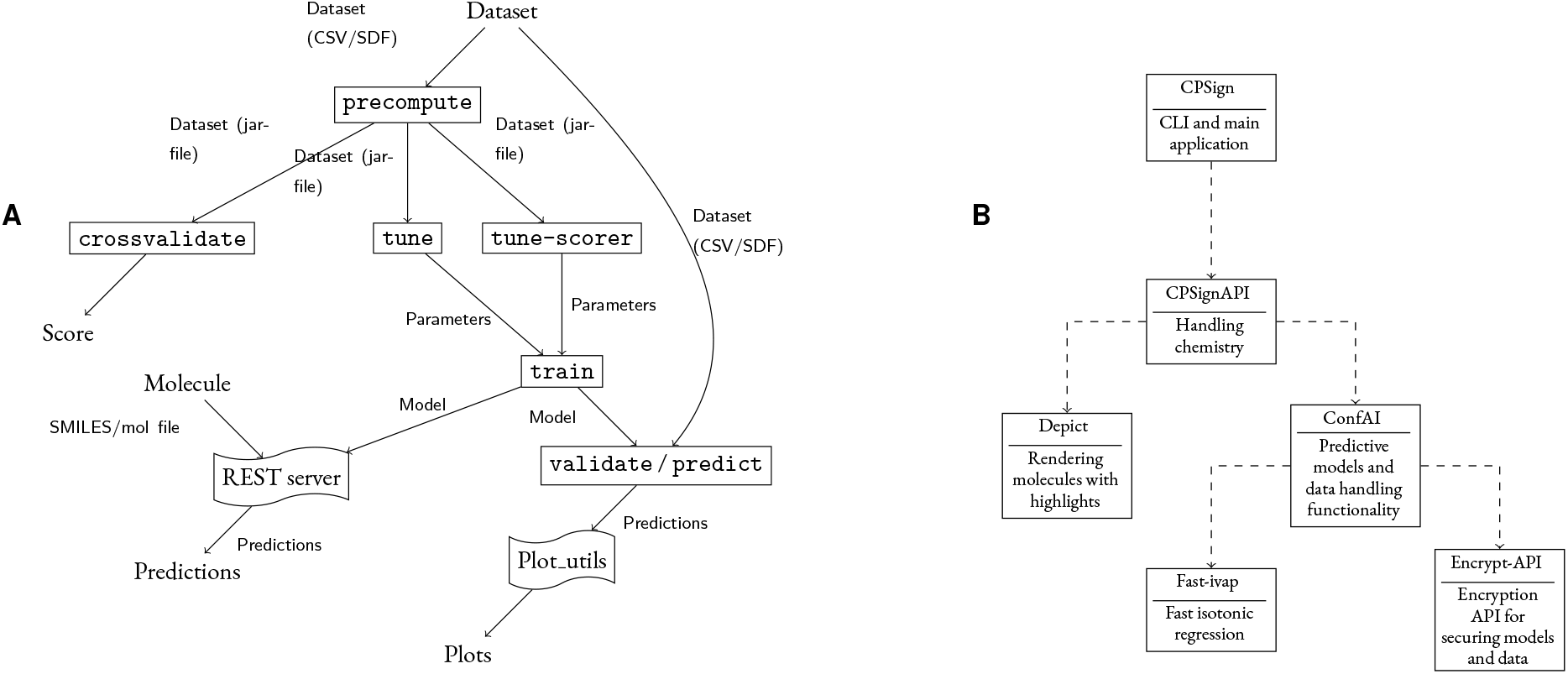
Panel A shows the general workflow of working with CPSign. Datasets are first precomputed then it is possible to do cross-validation or tune hyper-parameters, either by just looking at the scorer or at the entire probability framework, e.g., conformal prediction. After that a final model can be trained and validated or used for predictions, or a model can be trained immediately using the default values for hyper-parameters. The final model can be published as a separate REST server and plots can be made using the separate Plot_utils Python package. Panel B depicts an UML diagram over the individual modules of CPSign, where arrows depict dependence towards another module.

### Adding custom extensions

There are several ways that users can add their own custom extensions to CPSign, e.g., by providing pull requests on GitHub, cloning the repository or simply by extending the desired interface and exposing it as a Java Service Provider. CPSign loads extendable interfaces using the `ServiceLoader` class which makes it possible with minor effort to add custom code that can be used even through the CLI. For an example of how this can be achieved, see our CPSign-DL4J extension at GitHub (https://github.com/arosbio/cpsign-dl4j) which makes it possible to use deep learning models as underlying scoring models by wrapping the DeepLearning4J [48] package. This DL4J extension was evaluated and developed as part of a thesis work [55], resulting in predictive models performing on par with the SVM based models.

### Interfaces

There are multiple ways of running CPSign, each explained in more detail in the following subsections. User documentation is found on https://cpsign.readthedocs.io. An overview of how CPSign can be used and the typical workflows are outlined in Figure 5A, and are here described briefly. CPSign works with tabular type data, in either CSV format containing chemistry in SMILES format, or SDF files. The first step is always to convert the chemical input file(s) into numerical data, which is performed using one or several descriptors and termed precompute. The `precompute` step results in a precomputed dataset, containing both numerical data and all meta data from that step, so e.g. the same descriptors and data transformations are applied to any future test molecules. From precomputed data, the user can either run `crossvalidate` to evaluate the given data with a predictor-setting (i.e. conformal or Venn-ABERS model, including specific settings for scoring model, nonconformity function and any additional hyper-parameters that can be set), to quickly assess the expected performance for a new dataset and settings.

From a precomputed dataset a predictor model can be trained, with an optional intermediate step of hyper-parameter tuning using either tune (hyper-parameter tuning including all tuneable predictor-parameters) or tune-scorer (hyper-parameter tuning of the underlying scorer model only). The `train` step can thus be run either using default parameters (or manually set parameters) or using tuned parameters from the optional tuning step. The trained model can then be validated with an external validation-set or used to predict new compounds. For the final trained models, there is also the option to deploy them as micro services which can be deployed locally or publicly, allowing users to run query predictions using REST, this option is further described in the section REST API.

### Command Line Interface

The Command Line Interface (CLI) is the main way that CPSign is intended to be used, in a high abstraction level which facilitate rapid evaluation of new datasets and models. Apart from the user documentation online, the CLI tool has a rich user manual available directly in the terminal environment, as well as a help program (`explain`) that both provide detailed explanations about key arguments and lists available settings. The listing functionality is useful as CPSign can be extended with custom implementations and the documentation is generated dynamically depending on what is currently available, including listing of sub-parameters dynamically.

Working with the CLI follows the outline in Figure 5A, having a separate “program” for each rectangle in the figure. The goal has been to make the CLI as feature complete as possible, while balancing the level of complexity of the interface. In this spirit most parameters have been set to good default parameters, favoring less computational complexity (such as using a linear kernel SVM as default), but always making it possible to change settings for more elaborate alternatives. Most users thus prefer working with the CLI, e.g. for publications [56,12, 57, 58] as well as other unpublished work.

### Java API

For greater control of all available tweaks and handles, and for incorporating CPSign in other programs the Java API can be used. Here the user can also chose to depend on another sub-module of CPSign (Figure 5B) depending on their specific requirements. Coding examples can be found in the CPSign-example GitHub repository (https://github.com/ arosbio/cpsign-examples), to make it easier for new users to start coding against the API.

### REST API

To make it easy for users to make their developed models publicly available, the final models can be deployed as micro services and users can interact with them using REST. Each service is automatically documented using the OpenAPI 3.0 specification [59], and can optionally include a graphical user interface in which the user can draw or paste chemical structures and get the predictions back as well as atom contributions drawn using the method described in the section Interpretation of predictions. The web service implementation is freely available in the CPSign_predict_services GitHub repository (https://github.com/arosbio/cpsign_predict_services), and can thus be altered according to any further requirements, e.g., by adding user identification. Examples of these services running in production are the models serving the web page accessible at https://predgui.serve.scilifelab.se.

### Plot utils

When implementing CPSign a decision was made to not include any plotting functionality into the program itself but instead let the user create figures with their own favorite tool for this task. Creating figures through a CLI would both restrict the level of flexibility as well as clutter the API with too many parameters. However, to make it easy to quickly generate figures to analyze results we developed a Python library building on top of the popular matplotlib [60], as well as added functions to load results from CPSign. This library can be found in the Plot_utils GitHub repository (https://github.com/pharmbio/plot_utils) and was used for generating e.g. Figure 2 and 3. An example of how to generate Figure 3D can be found in Listing 1. Note that the regression case is more difficult compared to classification, as the confidence/significance level must be given at prediction time and the lower/upper bounds of each prediction interval must be saved and loaded. For classification the loading only requires picking the columns containing p-values from a CSV file, and significance levels can be applied when generating the figures.

**Listing 1:**
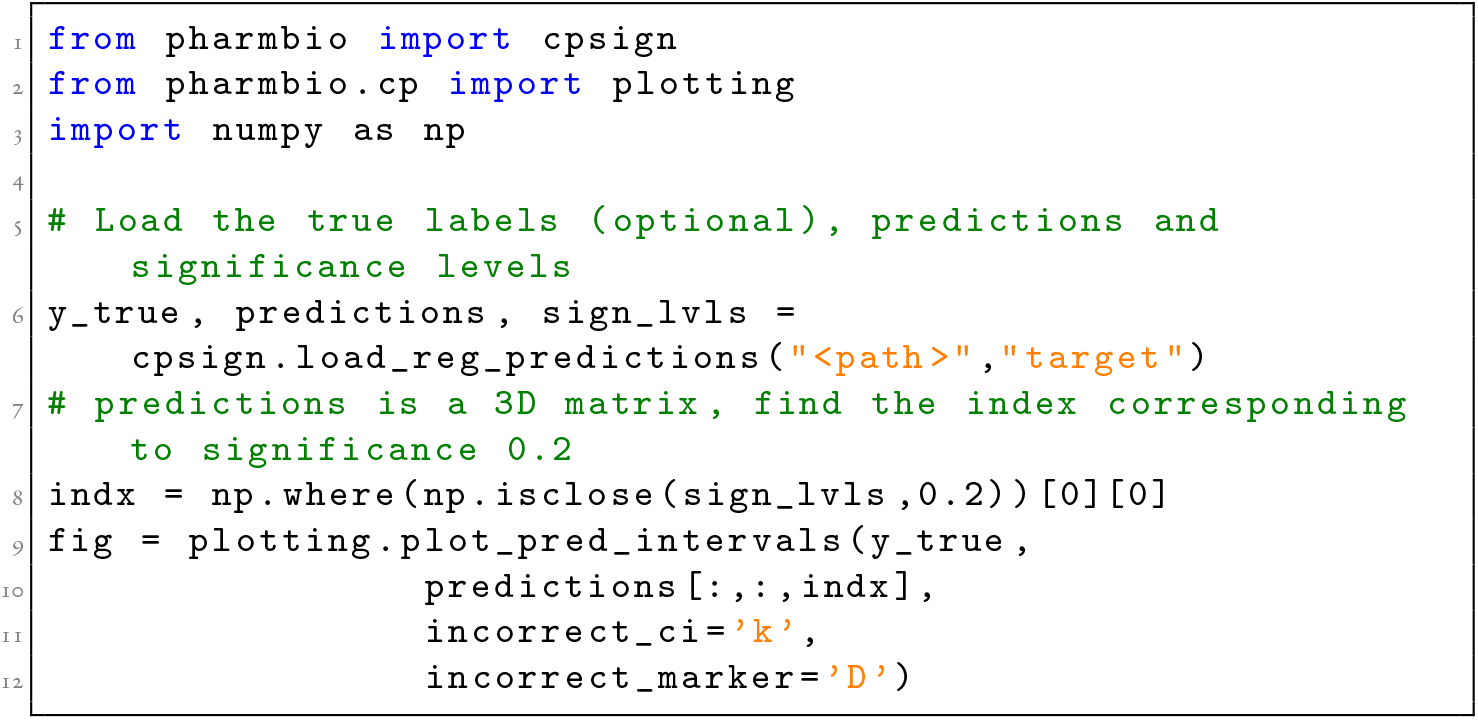
Loading predictions from a conformal regression model and plotting the predicted intervals at a given sigificance level (generating Figure 3D). Note that the predictions output is a 3D ndarray where a slice of the two first dimensions give the predictions at a single sigificance level.

## Evaluation

In this section we make a comparison of CPSign versus other common modeling approaches used for QSAR modeling, both the traditional method of handcrafted Morgan fingerprints combined with Random Forest, as well as the contemporary graph neural network based Chemprop [61]. The objective here is not to make a comprehensive comparison across all combinations of descriptors and modeling methods that are currently available, but rather using representative methods and data sets.

### Datasets

For the evaluation we use a subset of the benchmarking datasets from the popular MoleculeNet [62]. The classification datasets were picked from both of the categories Biophysics and Physiology in order to obtain a broader range of tasks. The selected datasets are outlined in Table 1. For regression the number of available datasets in MoleculeNet were limited, so all tasks that were not part of the category Quantum mechanics were selected, see Table 2. The reason for excluding the Quantum mechanics datasets was that they are based on crystal structures of protein-ligand complexes, for which CPSign lack descriptors for modeling. To expand the evaluation, the 13 largest curated datasets published in Skuta et al. [63] (additional file 1) were included, as well as the two largest datasets from Papyrus [64]. This resulted in 16 datasets for classification and 18 for regression.

**Table 1:** The 16 classification data sets used in the evaluation, taken from the MoleculeNet benchmark datasets. All splits into train, calibrate and test were performed stratified.

| Dataset | Sub-task | Train | Calibrate | Test | Class ratio<br>(inactive:active) |
| --- | --- | --- | --- | --- | --- |
| Tox21 | NR-AR | 4 359 | 1 453 | 1 453 | 6 956 : 309 |
|  | NR-AR-LBD | 4 054 | 1 352 | 1 352 | 6 521 : 237 |
|  | NR-AhR | 5 258 | 648 | 643 | 5 781 : 768 |
|  | NR-Aromatase | 3 492 | 1 164 | 1 165 | 5 521 : 300 |
|  | NR-ER | 4 949 | 618 | 626 | 5 400 : 793 |
|  | NR-ER-LBD | 4 173 | 1 391 | 1 391 | 6 605 : 350 |
|  | NR-PPAR-gamma | 3 870 | 1 290 | 1 290 | 6 264 : 186 |
|  | SR-ARE | 4 671 | 584 | 577 | 4 890 : 942 |
|  | SR-ATAD5 | 4 243 | 1 414 | 1,415 | 6 808 : 264 |
|  | SR-HSE | 3 880 | 1 293 | 1 294 | 6 095 : 372 |
|  | SR-MMP | 4 661 | 568 | 581 | 4 892 : 918 |
|  | SR-p53 | 4 064 | 1 355 | 1 355 | 6 351 : 423 |
| HIV | - | 32 901 | 4 113 | 4 113 | 39 684 : 1 443 |
| PCBA | PCBA-686978 | 241 714 | 30 316 | 30 145 | 239 375 : 62 800 |
|  | PCBA-884 | 8 037 | 1 005 | 1 005 | 6 719 : 3 328 |
|  | PCBA-914 | 4 537 | 1 513 | 1 513 | 7 346 : 217 |

**Table 2:**
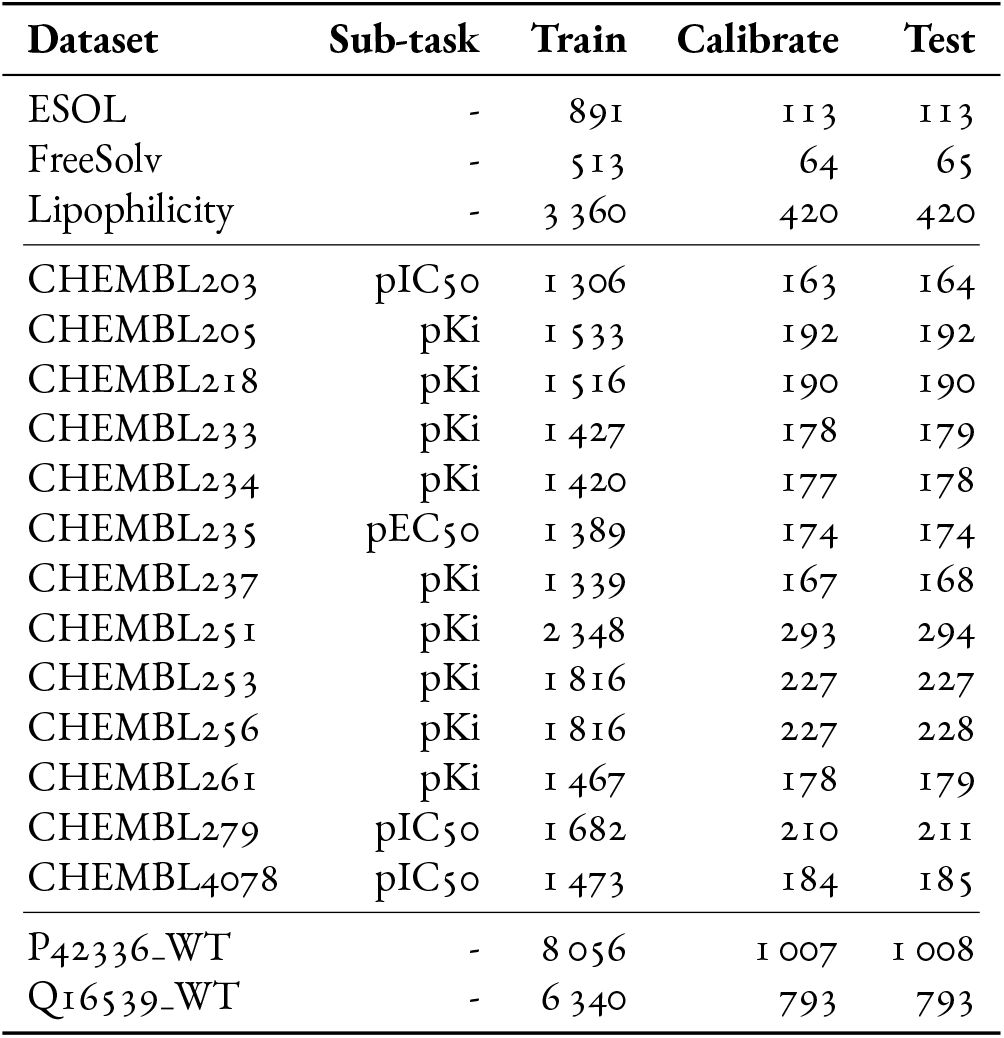
The 18 regression data sets used in the evaluation, were the three first datasets comes from the MoleculeNet benchmark datasets, the 13 in the middle from Skuta et al. and the remaining two from Papyrus. Splitting into train, calibrate and test sets were made randomly.

Each dataset was split into three subsets; `train`, `calibrate` and `test`, in ether 80 % - 10 % - 10 % splits, or 60 % - 20 % - 20 % splits for the classification datasets with fewer than 500 observations for the minority class (see Table 1 and 2). The datasets from MoleculeNet were downloaded and split using the deepchem software [65], the splitting was performed randomly for regression and random stratified for classification datasets. The datasets from Skuta et al. and Papyrus were split randomly using numpy [66].

### Modeling methods

In this comparison the Inductive Conformal Predictor (ICP) was used by all methods, with predefined splits of *proper training* and *calibration sets* according to the train and calibrate splits described in the previous paragraph - so all methods had exactly the same training, calibration and testing data. The discerning factors between the methods was the descriptors, the underlying scoring models, nonconformity function and any additional parameters that can be tweaked within each software, such as employing interpolation of the p-values to smooth out the predictions. The following modeling methods were used in the comparison:

#### CPSign

CPSign using the CLI with the default descriptor, i.e. the Signatures descriptor [27, 28], with an RBF-kernel SVM except for the largest classification dataset (PCBA-686978) for which a linear kernel SVM was used instead. The default nonconformity function was used for both classification (negative distance to SVM hyperplane) and regression (LogNormalized). The LogNormalized function uses an additional error model to estimate the difficulty of each example, which is used to scale the prediction interval. Additionally, linear interpolation was employed for the p-value calculation. All other settings were the default ones.

#### CPSign tuned

This strategy used the same parameters as described for CPSign above, but extended with a hyper-parameter tuning step of the SVM hyper-parameters cost (*C*) and gamma (*γ*) using grid search. The grid consisted of ten values for *C* (2^−6^, 2^−4^,…, 2^12^) and six for *y* (2^−14^, 2^−12^,…, 2^−4^) for a total of 60 combinations, apart from the largest classification dataset which used the linear kernel SVM where only the ten *C* values were evaluated. Hyper-parameter tuning was exclusively performed on the train partition of the data, running a 10-fold cross-validation of the SVM only (i.e. without adding the conformal calibration) and optimizing with respect to macro Fi score for classification and RMSE for regression.

#### FP+RF tuned

Random Forest (RF) using Morgan Fingerprints (FP) as descriptor and nonconformist [21] for CP implementation. The Morgan Fingerprints were calculated using RDKit [67], using bit length of 2048 and radius 2. The RF hyper-parameters were tuned using the `train` split, without including conformal calibration, optimizing for balanced accuracy for the classification datasets and RMSE for the regression datasets. The grid of tested hyper-parameters had 32 combinations for the classification models and 64 for the regression models. For the conformal classification model the `MarginErrFunc` was used as nonconformity function and for regression the AbsErrFunc was used in conjunction with a normalizer model, using an additional RF model using the same hyper-parameters as the scoring model.

#### Chemprop

The Chemprop software [68, 61] was used to develop Directed Message Passing Neural Network (D-MPNN) models. Default parameter settings and network architecture was used. A separate validation set (10 %) was randomly split off from the train dataset for monitoring model training (using random stratified splitting for classification). For the classification models 1-probability for the class was used as nonconformity measure, and using the smoothed calculation of p-values (i.e. special treatment of equal nonconformity scores). The procedure described in Norinder et al. [29] was used for regression, i.e., using one model to predict the midpoint and a second (error-model) for predicting the error made by the first model in order to normalize the prediction intervals based on the predicted difficulty of the object. The nonconformity function was extended by adding a small smoothing factor, *β*, of 0.01 as CPSign does for the normalized nonconformity measures (which increases stability as well as removes the potential of division by 0 in the calculation).

#### Chemprop tuned

This method used the same settings as the method above, but with the added step of hyper-parameter tuning of the Chemprop model using the `chemprop_hyperopt` function. Chemprop performs a bayesian hyper-parameter optimization using the hyperopt package [69], here evaluated using the default 20 different hyper-parameter settings. To minimize information leakage, this optimization step was only applied on the 90 % split from the train dataset, and thus chemprop internally split that set further into a validation set for monitoring the model training, a test-set to compare the model performance of different hyper-parameters and data used for training the model. For the regression experiments the error model used the same optimized hyper-parameters as the scoring model that predicted the midpoint.

### Comparison

While comparing the methods we restricted the analyzed significance levels to 0.01-0.3 for classification and 0.05-0.3 for regression, corresponding to 70-99 and 70-95 % confidence. The methods were first assessed with respect to calibration using calibration plots, shown in Figure S2 and S3. To simplify and quantify the calibration of the different methods we computed the maximum (signed) difference between the error rate and specified significance level, the RMSE of error rate against significance level as well as the “capped” RMSE, Figure S1. The capped RMSE was calculated by setting the error rate equal to the significance level if it was lower than the significance level (for every evaluated significance level), so that over-conservative predictions (i.e. lower error rate than required) do not contribute to a higher RMSE. The guarantees made by the conformal framework is that the error rate should be at most equal to the significance level, the capped RMSE is more representative of level of calibration.

All methods produce similar results with respect to calibration, where the only concern is the calibration of the minority class for some of the classification datasets. This is most likely due to the smaller number of observations of the minority class in both the calibration and test splits, which leads to higher variance. The methods perform similar enough so they can be compared fairly with respect of predictive efficiency.

### Classification

Aggregating the predictive efficiency across the datasets resulted in similar predictive efficiency for all evaluated methods (Figure 6A-B). Differences between methods can be seen when analyzing the datasets individually in terms of Observed Fuzziness (Figure S4) and fraction of single-label predictions (Figure S5), where the most notable trend were that the two Chemprop methods preformed best on the two largest datasets (HIV and PCBA-686978). The results were further ranked (Table S2) to find the top-performer as well as their overall rank, where the CPSign method (i.e., without tuning) was the overall best method.

**Figure 6:**
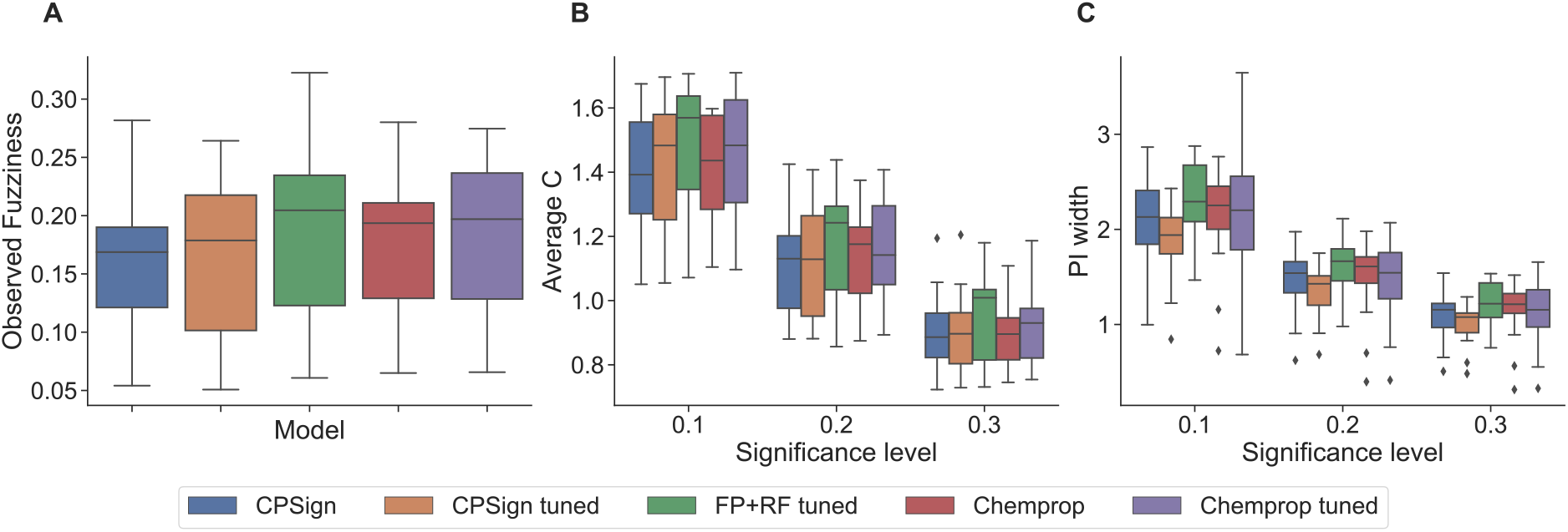
Boxplots for aggregating the results from all tested datasets from the evaluation, panel A-B are from the classification experiments, panel C for the regression experiments. Panel A; the Observed Fuzziness (lower values are preferable), this metric is independent of significance level. Panel B; average number of predicted classes (Average C), lower values are preferable. Panel C; Prediction Interval (PI) width for the regression experiments, lower values are preferable. Model performance is very similar across all tested methods, where there is only a discernible difference in panel C where the CPSign tuned box is visually lower than the rest - but without statistical significance using a nonparametric Wilcoxon signed-rank test. Notably the whiskers for the Chemprop tuned method in panel C show that its prediction intervals varies more compared to the other methods (both positively and negatively).

### Regression

Aggregating the results across all evaluated datasets (Figure 6C) shows that the CPSign tuned method generated the most efficient predictions overall, although the methods again performs largely similarly. Calculating the Wilcoxon signed-rank test between each combination of method, separately for each significance level, gave no significant difference between any method. All results presented separately for each dataset can be found in Figure S6, and the rankings across all datasets in Table S2. From the ranking it is clear that the CPSign tuned method performed best overall, but that each method was the top-performing method for at least one dataset and significance level.

### Runtime comparison

Comparison was further performed in respect to runtime for each experiment to complete. Both CPSign methods and the FP+RF tuned method were run on a laptop whereas the Chemprop experiments were run on computer cluster with a Nvidia 1080 Ti GPU. A summary of the runtimes can be found in Table 3, with individual datasets in supplemental Figures S7 and S8. Note that no replicate runs were performed, so the results should be interpreted as an indication of how the methods compare.

**Table 3:** Runtime comparison summarized across all datasets, were each runtime is calculated to be relative to the CPSign method as a baseline and the reported values are the median over all datasets. The median value was chosen as the experimental setups varied, e.g., as CPSign was run with different kernels, both directly affecting training time for the SVM, and the size of the parameter search grid, leading to large fluctuations depending on which dataset is considered. The value 59 x for CPSign tuned e.g. specify that it took 59 times as long for CPSign tuned to complete compared to CPSign to complete.

| Model strategy | Runtime relative baseline |  |
| --- | --- | --- |
|  | Classification | Regression |
| CPSign | 1 | 1 |
| CPSign tuned | 59 x | 32 x |
| FP+RF tuned | 6.0 x | 43 x |
| Chemprop | 28 x | 38 x |
| Chemprop tuned | 510 x | 350 x |

## Discussion

When making predictions on novel chemical structures and where it is certain that it has not been used in model training, scientists face the inevitable questions of how much to trust the prediction, and if trusted then how to interpret the result. While most traditional methods provide a single level of model confidence from the training procedure, e.g. a metric from cross-validatation, CPSign via the conformal prediction methodology outputs a prediction interval that is specific for each predicted object. If the object to predict is more different to what has been seen before, the prediction interval becomes larger, and the user can be sure to trust the size of these intervals as they are based on proven mathematical theory [6]. This also offers a compelling alternative to the concept of Applicability Domain [5]. It is important to notice that the size of prediction intervals are related to the choice of nonconformity function; a poor nonconformity function will still be valid but result in larger prediction intervals [70, 71]. Since conformal prediction outputs a prediction interval given a user-specified level of confidence, it can be difficult to choose this level of confidence; if requiring a higher confidence then the prediction intervals will naturally be larger. In the end this comes down to the user having to make a decision on what is acceptable for the specific problem given the interval sizes produced by the model. In the author’s experience this can lead to a more realistic view on model expectations.

Recently, Deep Neural Networks (DNNs) have emerged as a popular method for supervised learning, and shown higher accuracy for many traditional machine learning tasks. A prominent example is computer vision when the objects are images and where convolutional neural networks (CNNs) have yielded dramatic improvements [72]. For tabular data the improvements are not as profound. DNNs generally require larger training sets compared to traditional machine learning methods, although techniques such as transfer learning and augmentation can somewhat reduce this burden [73]. Further, DNNs necessitate hyper-parameter tuning on a much larger scale than traditional machine learning methods, making them costly in terms of time and computational resources.

For supervised learning where the data objects are chemical structures, several studies have been presented to compare different deep learning approaches with more traditional machine learning methods, such as [74, 75]. One problem with comparisons lies in the choice of metric, for example using accuracy or AUC (AUROC) is not suitable when working with unbalanced data that is very common in the field. Another and more serious problem is that studies rarely assess the level of calibration of models. Deep Learning models have been shown in many cases to be poorly calibrated [76], rendering comparisons on the produced output probabilities biased. Conformal prediction is one method to calibrate models to obtain valid (well-calibrated) probability estimates, and in our evaluation we use it compare CPSign with DNNs as implemented in the Chemprop package both in terms of calibration and efficiency. As conformal prediction produces prediction intervals, traditional metrics such as AUC and F1 cannot be used, and we instead use the well-established metrics of Observed Fuzziness, Average C and fraction of single-label prediction intervals. Using mondrian conformal prediction [6] also improves the modeling and calibration for imbalanced datasets, which has been shown in several ligand-based studies [30,18, 31].

Comparing CPSign against other popular modeling methods showed that overall all methods performed similarly in terms of predictive efficiency for the classification datasets. For the regression datasets CPSign with tuned hyper-parameters was overall the top-performing method (Figure 6). Looking at individual datasets each modeling method was the top-performing method at least once, showing that the optimal modeling approach can vary depending on the data being modeled. We note for instance that the DNN-based Chemprop performed better on the largest datasets, supporting the hypothesis that DNN requires large training sets. Our overall conclusion is that CPSign performs on par with DNNs (Chemprop), when calibrating models using conformal prediction and hence take advantage of theoretically proven model validity, which also was assessed empirically.

The lack of interpretability of DNNs is widely acknowledged [77]. CPSign utilizes the Signatures descriptors to represent chemistry, which allows for feature importance to be visualized as chemical substructures. Due to the fast predictions, CPSign allows for immediate feedback and visualization of atom contributions (Figure 4) to the prediction (“calculate-as-you-draw”). This has been a much appreciated feature by users of the software, such as medicinal chemists.

Setting up and maintaining computational environments for machine learning can be demanding, and this is especially evident for deep neural networks having many and specific dependencies. Further, with changing versions of frameworks and dependencies, it can be time-consuming to maintain models and predictions over time, such as in production environments [78]. The requirements on IT infrastructure also varies a lot between modeling methods, with DNNs generally requiring access to GPUs to accelerate the learning. Even so, the amount of hyper-parameters that needs to be optimized commonly lead to several days of model training. In contrast, CPSign has a single dependency (Java) that makes it straightforward to download, use, integrate into other systems, and with a modeling that is fast to complete. Examples of tools and systems based on CPSign are ANDROMEDA by Prosilico (https://prosilico.com/andromeda) and PredGUI in SciLifeLab Serve (https://predgui.serve.scilifelab.se).

CPSign is an advanced tool with many options, and hence it requires abit of learning to understand the parameters of the software. Effort has been made to reduce the number of required parameters, and to provide good default values. For example, the default option of CPSign is Inductive Conformal Prediction (ICP) requiring a number of data points to be set aside into a calibration set. This is the price to pay for obtaining valid (well-calibrated) models. When having few data points in the training set, there is always the option to use Transductive Conformal Prediction (TCP) that does not use a calibration set, but with the downside that each prediction requires a re-training of the model. However when data sizes are so low that TCP is mandated, this is usually an acceptable tradeoff.

For SVM, it generally leads to more efficient models when using a non-linear kernel such as RBF. However, this is computationally expensive when datasets are large, although it has been shown that for larger models the difference between linear and non-linear kernels is small [46]. The default setting for CPSign is to use a linear kernel (LIBLINEAR) to produce fast results when prototyping, and it is recommended to switch to RBF kernel if datasets are of small or moderate size, also depending on the computational infrastructure available. As the comparison shows (Figure 6, S4, S5, S6), the default values for parameters *C* and *γ* as previously devised [47] are generally well performing, but in some cases the efficiency can be improved using hyper-parameter tuning.

Although being a worthy tool for many typical cheminformatics modeling tasks, it is worth mentioning that there are tasks that CPSign is not suitable for, such as for non-tabular data (e.g. images or graph-based data) or when multi-task learning could be employed.

## Conclusion

CPSign is a robust and complete implementation of conformal prediction for cheminformatics applications. The combination of signatures and SVM has been shown to produce robust and accurate models, and conformal prediction adds the ability to produce valid prediction intervals. The implementation as a single software package with no dependencies apart from a Java runtime, a well-developed API, and a low footprint makes it suitable both for rapid prototyping and integration in production system. The evaluation of modeling methods highlights that CPSign performs overall on par or outperforms other state-of-the-art cheminformatics approaches.

## Acknowledgments

OS acknowledges funding from the Swedish Research Council (grants 2020-03731 and 2020-01865), FORMAS (grant 2022-00940), Swedish Cancer Foundation (22 2412), and Horizon Europe grant agreement #101057014 (PARC).

## References

[1] Jessica Vamathevan et al. “Applications of machine learning in drug discovery and development”. en. In: Nat. Rev. Drug Discov. 18.6 (June 2019), pp. 463–477.

[2] Anna O Basile, Alexandre Yahi, and Nicholas P Tatonetti. “Artificial Intelligence for Drug Toxicity and Safety”. en. In: Trends Pharmacol. Sci. 40.9 (Sept. 2019), pp. 624–35.

[3] Eugene N Muratov et al. “QSAR without borders”. en. In: Chem. Soc. Rev. 49.11 (June 2020), pp. 3525–3564.

[4] Jose Jimenez-Luna et al. “Artificial intelligence in drug discovery: recent advances and future perspectives”. en. In: Expert Opin. Drug Discov. 16.9 (Sept. 2021), pp. 949–959.

[5] Domenico Gadaleta et al. “Applicability domain for QSAR models: where theory meets reality”. In: International journal of quantitative structure-property relationships (IJQSPR) 1.1 (2016), pp. 45–63.

[6] Vladimir Vovk, Alex Gammerman, and Glenn Shafer. Algorithmic Learning in a Random World. New York: Springer, Jan. 2005. doi: 10.1007/b106715.

[7] Ulf Norinder et al. “Introducing conformal prediction in predictive modeling. A transparent and flexible alternative to applicability domain determination”. In: J Chem Inf Model 54.6 (June 2014), pp. 1596–603. doi: 10.1021/ci5001168.

[8] U Norinder, A Rybacka, and P L Andersson. “Conformal prediction to define applicability domain - A case study on predicting ER and AR binding”. In: SAR QSAR Environ Res 27.4 (Apr. 2016), pp. 303–16. doi: 10.1080/1062936X.2016.1172665.

[9] Jonathan Alvarsson et al. “Predicting with confidence: using conformal prediction in drug discovery”. In: Journal of Pharmaceutical Sciences 110.1 (2021), pp. 42–49.

[10] Fredrik Svensson et al. “Maximizing gain in high-throughput screening using conformal prediction”. In: J Cheminform 10.1 (Feb. 2018), p. 7. doi: 10.1186/s13321-018-0260-4.

[11] Fredrik Svensson, Ulf Norinder, and Andreas Bender. “Modelling compound cytotoxicity using conformal prediction and PubChem HTS data”. In: Toxicol Res (Camb) 6.1 (Jan. 2017), pp. 73–80. doi: 10.1039/c6tx00252h.

[12] Andrea Morger et al. “Assessing the calibration in toxicological in vitro models with conformal prediction”. In: Journal of Cheminformatics 13.1 (2021), p. 35.

[13] Maris Lapins et al. “A confidence predictor for logD using conformal regression and a support-vector machine”. In: J Cheminform 10.1 (Apr. 2018), p. 17. doi: 10.1186/s13321-018-0271-1.

[14] Samuel Lampa et al. “Predicting Off-Target Binding Profiles With Confidence Using Conformal Prediction”. In: Front Pharmacol 9 (2018), p. 1256. doi: 10.3389/fphar.2018.01256.

[15] Urban Fagerholm et al. “In silico predictions of the human pharmacokinetics/toxicokinetics of 65 chemicals from various classes using conformal prediction methodology”. In: Xenobiotica 52.2 (Feb. 2022), pp. 113–118. doi: 10.1080/00498254.2022.2049397.

[16] Urban Fagerholm et al. “In Silico Prediction of Human Clinical Pharmacokinetics with ANDROMEDA by Prosilico: Predictions for an Established Benchmarking Data Set, a Modern Small Drug Data Set, and a Comparison with Laboratory Methods”. In: Altern Lab Anim 51.1 (Jan. 2023), pp. 39–54.doi: 10.1177/02611929221148447.

[17] Isidro Cortes-Ciriano and Andreas Bender. “Deep Confidence: A Computationally Efficient Framework for Calculating Reliable Prediction Errors for Deep Neural Networks”. In: J Chem Inf Model 59.3 (Mar. 2019), pp. 1269–1281. doi: 10.1021/acs.jcim.8b00542.

[18] Ulf Norinder. “Traditional Machine and Deep Learning for Predicting Toxicity Endpoints”. In: Molecules 28.1 (Dec. 2022). doi: 10.3390/molecules28010217.

[19] Jin Zhang, Ulf Norinder, and Fredrik Svensson. “Deep Learning-Based Conformal Prediction of Toxicity”. In: J Chem Inf Model 61.6 (June 2021), pp. 2648–2657. doi: 10.1021/acs.jcim.1c00208.

[20] Henrik Olsson et al. “Estimating diagnostic uncertainty in artificial intelligence assisted pathology using conformal prediction”. In: Nat Commun 13.1 (Dec. 2022), p. 7761. doi: 10.1038/s41467-022-34945-8.

[21] Henrik Linusson. Nonconformist. 2015. URL: http://donlnz.github.io/nonconformist/.

[22] Nicolas Bosc et al. “Large scale comparison of QSAR and conformal prediction methods and their applications in drug discovery”. In: Journal of cheminformatics 11 (2019), pp. 1–16.

[23] Fredrik Svensson, Ulf Norinder, and Andreas Bender. “Improving screening efficiency through iterative screening using docking and conformal prediction”. In: Journal of chemical information and modeling 57.3 (2017), pp. 439–444.

[24] U Norinder et al. “Conformal prediction of HDAC inhibitors”. In: SAR QSAR Environ Res 30.4 (Apr. 2019), pp. 265–277. doi: 10.1080/1062936X.2019.1591503.

[25] Henrik Bostrom. “crepes: a Python Package for Generating Conformal Regressors and Predictive Systems”. In: Conformal and Probabilistic Prediction with Applications. PMLR. 2022, pp. 24–41.

[26] Valery Manokhin. Awesome Conformal Prediction. Version v1.0.0. Apr. 2022. doi: 10.5281/zenodo.6467205. url: 10.5281/zenodo.6467205.

[27] Jean-Loup Faulon, Donald P. Visco, and Ramdas S. Pophale. “The Signature Molecular Descriptor. 1. Using Extended Valence Sequences in QSAR and QSPR Studies”. In: J. Chem. Inf. Model. 43.3 (2003). PMID: 12767129, pp. 707–720. doi: 10.1021/ci020345w. eprint: https://doi.org/10.1021/ci020345w. url: https://doi.org/10.1021/ci020345w.

[28] Jean-Loup Faulon, Carla J Churchwell, and Donald P Visco. “The signature molecular descriptor. 2. Enumerating molecules from their extended valence sequences”. In: J. Chem. Inf. Model. 43.3 (2003). PMID: 12767130, pp. 721–734. doi: 10.1021/ci020346o.

[29] Ulf Norinder, et al. “Introducing conformal prediction in predictive modeling. A transparent and flexible alternative to applicability domain determination”. In: J. Chem. Inf. Model. 54.6 (2014), pp. 1596–1603.

[30] Jiangming Sun et al. “Applying Mondrian Cross-Conformal Prediction To Estimate Prediction Confidence on Large Imbalanced Bioactivity Data Sets”. In: J. Chem. Inf. Model. 57.7 (July 2017), pp. 1591–1598. doi: 10.1021/acs.jcim.7b00159.

[31] Ulf Norinder and Scott Boyer. “Binary classification of imbalanced datasets using conformal prediction”. In: J. Mol. Graphics Modell. 72 (2017), pp. 256–265.

[32] Vladimir Vovk et al. “Criteria of efficiency for conformal prediction”. In: Symp. on Conformal and Probabilistic Prediction with Appl. Springer. 2016, pp. 23–39.

[33] Vladimir Vovk. “Venn predictors and isotonic regression”. In: CoRR abs/1211.0025 (2012).

[34] Vladimir Vovk, Ivan Petej, and Valentina Fedorova. “Large-scale probabilistic prediction with and without validity guarantees”. In: Proceedings of NIPS. Vol. 2015. 2015.

[35] Dirar Sweidan and Ulf Johansson. “Probabilistic Prediction in scikit-learn”. In: The 18th International Conference on Modeling Decisions for Artificial Intelligence, Online (from Umea, Sweden), September 27-30, 2021. 2021.

[36] Ruben Buendia et al. “Accurate hit estimation for iterative screening using venn-abers predictors”. In: Journal of Chemical Information and Modeling 59.3 (2019), pp. 1230–1237.

[37] Staffan Arvidsson et al. “Prediction of Metabolic Transformations using Cross Venn-ABERS Predictors”. In: Conformal and Probabilistic Prediction and Applications. PMLR. 2017, pp. 118–131.

[38] Ernst Ahlberg, Ruben Buendia, and Lars Carlsson. “Using Venn-Abers predictors to assess cardio-vascular risk”. In: Conformal and Probabilistic Prediction and Applications. PMLR. 2018, pp. 132–146.

[39] Roberto Todeschini and Viviana Consonni. Handbook of molecular descriptors. John Wiley & Sons, 2008.

[40] Robert C Glen, et al. “Circular fingerprints: flexible molecular descriptors with applications from physical chemistry to ADME”. In: IDrugs 9.3 (2006), p. 199.

[41] Harry L Morgan. “The generation of a unique machine description for chemical structures-a technique developed at chemical abstracts service.” In: Journal of chemical documentation 5.2 (1965), pp. 107–113.

[42] David Rogers and Mathew Hahn. “Extended-connectivity fingerprints”. In: Journal of chemical information and modeling 50.5 (2010), pp. 742–754.

[43] Christoph Steinbeck, et al. “Recent developments of the chemistry development kit (CDK)-an open-source java library for chemo-and bioinformatics”. In: Current pharmaceutical design 12.17 (2006), pp. 2111–2120.

[44] Rong-En Fan, et al. “LIBLINEAR: A Library for Large Linear Classification”. In: J. of Machine Learning Research 9 (2008), pp. 1871–1874.

[45] Chih-Chung Chang and Chih-Jen Lin. “LIBSVM: A library for support vector machines”. In: ACM Transactions on Intelligent Systems and Technology 2 (3 2011). Software available at http://www.csie.ntu.edu.tw/~cjlin/libsvm, 27:1–27:27.

[46] Jonathan Alvarsson et al. “Large-scale ligand-based predictive modelling using support vector machines”. In: Journal of Cheminformatics 8.1 (2016), pp. 1–9.

[47] Jonathan Alvarsson, et al. “Benchmarking study of parameter variation when using signature fingerprints together with support vector machines”. In: J. Chem. Inf. Model. 54.11 (2014), pp. 3211–3217.

[48] Eclipse Deeplearning4j Development Team. Detplearning4j: Open-source distributed deep learning for the JVM. 2023. url: https://deeplearning4j.konduit.ai/.

[49] Lars Carlsson, Martin Eklund, and Ulf Norinder. “Aggregated Conformal Prediction”. In: Artif. Intell. Appl. and Innov. Ed. by Lazaros Iliadis et al. IFIPAICT 14. Berlin, Heidelberg: Springer Berlin Heidelberg, 2014, pp. 231–240. isbn: 978-3662-44722-2.

[50] Vladimir Vovk. “Cross-Conformal Predictors”. In: Ann. Math. Artif. Intell. 74.1-2 (June 2015), pp. 9–28. issn: 1012-2443. doi: 10.1007/s10472-013-9368-4. url: https://doi.org/10.1007/s10472-013-9368-4.

[51] Staffan Arvidsson McShane et al. “Machine learning strategies when transitioning between biological assays”. In: Journal of Chemical Information and Modeling 61.7 (2021), pp. 3722–3733.

[52] Ulf Johansson et al. “Handling small calibration sets in mondrian inductive conformal regressors”. In: Int. Symp. on Statistical Learning and Data Sci. Springer. 2015, pp. 271–280.

[53] Lars Carlsson et al. “Modifications to p-values of conformal predictors”. In: Int. Symp. on Statistical Learning and Data Sci. Springer. 2015, pp. 251–259.

[54] Ernst Ahlberg et al. “Interpretation of conformal prediction classification models”. In: Statistical Learning and Data Sciences: Third International Symposium, SLDS 2014, Egham, UK, April 20-23, 2013, Proceedings 3. Springer. 2015, pp. 323–334.

[55] Maria Deligianni. Comparison of Support Vector Machines and Deep Learning For QSAR with Conformal Prediction. 2022.

[56] Urban Fagerholm et al. “In silico prediction of human clinical pharmacokinetics with ANDROMEDA by Prosilico: Predictions for an established benchmarking data set, a modern small drug data set, and a comparison with laboratory methods”. In: Alternatives to Laboratory Animals (2023), p. 02611929221148447.

[57] Samuel Lampa, et al. “Predicting *off-target* binding profiles with confidence using conformal prediction”. In: Frontiers in Pharmacology 9 (2018), p. 1256.

[58] Maris Lapins et al. “A confidence predictor for logD using conformal regression and a support-vector machine”. In: Journal of cheminformatics 10 (2018), pp. 1–10.

[59] SmartBear Software. OpenAPI Specification. 2023. URL: https://swagger.io/specification/.

[60] J. D. Hunter. “Matplotlib: A 2D graphics environment”. In: Computing in Science & Engineering 9.3 (2007), pp. 90–95. doi: 10.1109/MCSE.2007.55.

[61] Esther Heid, et al. “Chemprop: Machine Learning Package for Chemical Property Prediction”. In: (2023).

[62] Zhenqin Wu, et al. “MoleculeNet: a benchmark for molecular machine learning”. In: Chemical science 9.2 (2018), pp. 513–530.

[63] C Skuta et al. QSAR-derived affinity fingerprints (part 1): fingerprint construction and modeling performance for similarity searching, bioactivity classification and scaffold hopping”. In: Journal of Cheminformatics 12.1 (2020), pp. 1–16.

[64] Olivier JM Bequignon et al. “Papyrus: a large-scale curated dataset aimed at bioactivity predictions”. In: Journal of cheminformatics 15.1 (2023), pp. 1–11.

[65] Bharath Ramsundar et al. Deep Learning for the Life Sciences. https://www.amazon.com/Deep-Learning-Life-Sciences-Microscopy/dp/1492039837. O’Reilly Media, 2019.

[66] Charles R. Harris et al. “Array programming with NumPy”. In: Nature 585.7825 (Sept. 2020), pp. 357–362. doi: 10.1038/s41586-020-2649-2. url: https:// doi.org/10.1038/s41586-020-2649-2.

[67] RDKit. *RDKit: Open-Source Cheminformatics Software*. URL: https://zenodo.org/record/7671152#.ZFIV43ZBzao.

[68] Kevin Yang et al. “Analyzing learned molecular representations for property prediction”. In: Journal of chemical information and modeling 59.8 (2019), pp. 33703388.

[69] James Bergstra, Daniel Yamins, and David Cox. “Making a science of model search: Hyperparameter optimization in hundreds of dimensions for vision architectures”. In: International conference on machine learning. PMLR. 2013, pp. 115–123.

[70] Martin Eklund, et al. “The application of conformal prediction to the drug discovery process”. In: Ann Math Artif Intell 74.1-2 (2015), pp. 117–132.

[71] Fredrik Svensson, et al. “Conformal regression for quantitative structure-activity relationship modeling—quantifying prediction uncertainty”. In: J. Chem. Inf. Model. 58.5 (2018), pp. 1132–1140.

[72] Alex Krizhevsky, Ilya Sutskever, and Geoffrey E Hinton. “Imagenet classification with deep convolutional neural networks”. In: Advances in neural information processing systems 25 (2012).

[73] Alexander Kensert, Philip J Harrison, and Ola Spjuth. “Transfer learning with deep convolutional neural networks for classifying cellular morphological changes”. In: SLAS Discovery: Advancing Life Sciences R&D 24.4 (2019), pp. 466–475.

[74] Zhenxing Wu, et al. “Do we need different machine learning algorithms for QSAR modeling? A comprehensive assessment of 16 machine learning algorithms on 14 QSAR data sets”. In: Briefings in bioinformatics 22.4 (2021), bbaa321.

[75] Alexandru Korotcov, et al. “Comparison of deep learning with multiple machine learning methods and metrics using diverse drug discovery data sets”. In: Molecular pharmaceutics 14.12 (2017), pp. 4462–4475.

[76] Chuan Guo et al. “On calibration of modern neural networks”. In: International conference on machine learning. PMLR. 2017, pp. 1321–1330.

[77] Igor I Baskin. “The power of deep learning to ligand-based novel drug discovery”. In: Expert opinion on drug discovery 15.7 (2020), pp. 755–764.

[78] Ola Spjuth, Jens Frid, and Andreas Hellander. “The machine learning life cycle and the cloud: implications for drug discovery”. In: Expert opinion on drug discovery 16.9 (2021), pp. 1071–1079.

