## Supplemental information for "CPSign - Conformal Prediction for Cheminformatics Modeling"

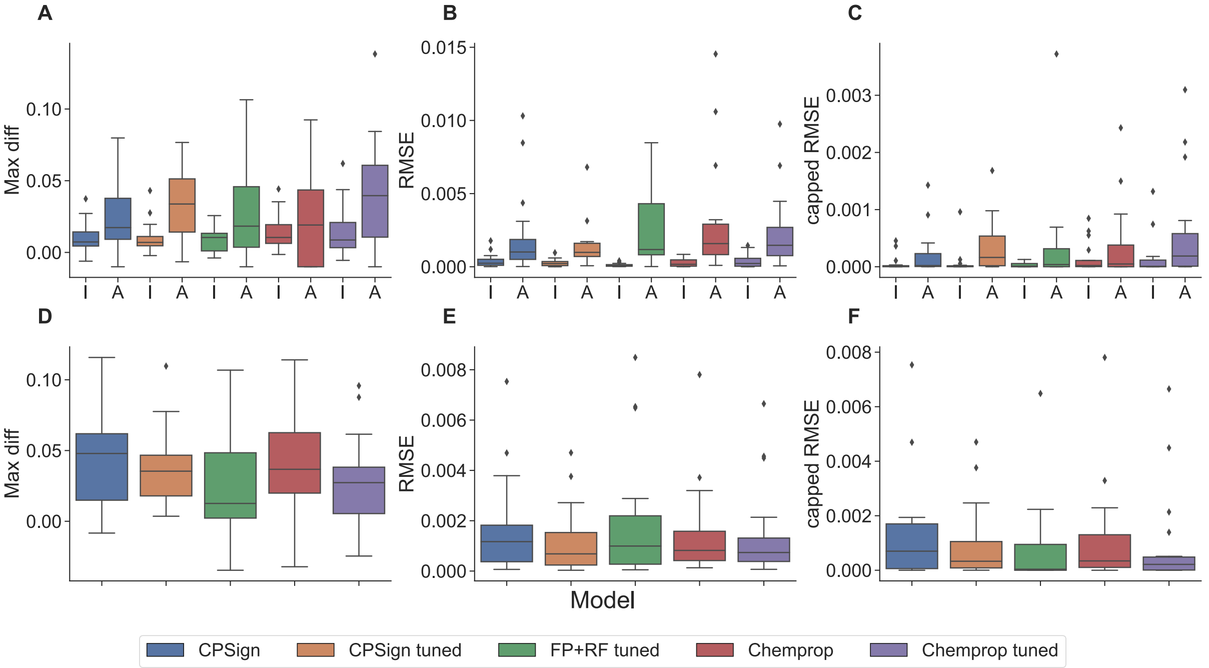


**Figure S1:** Calibration results aggregated for all evaluated methods and datasets, panels A-C for classification models, and panels D-F for regression models. For classification the calibration is analyzed independently per class (I: inactive, A: active), where the active class is the minority class for all datasets (Table 1). The classification values were based on 30 significance levels (0.01,0.02,...,0.3) whereas the regression ones were based on six levels (0.05,0.1,...,0.3). Panel A and B display "max diff" corresponding to the expression $\max_{\varepsilon}\{error\_rate_{\varepsilon} - \varepsilon\}$, i.e. the signed difference of error-rate and significance level, where a negative value corresponds to the error rate being smaller than the significance level across all tested significance levels, and a positive value means that the error rate exceeded the significance level with at most that difference (smaller values are preferable). Panel B and E display the root mean squared error (RMSE) between the significance level and the error rate (smaller values are preferable). Panels C and F display the "capped" RMSE, in which the error rate is capped at the significance level if it is lower than the significance level (for every evaluated significance level), so that over-conservative predictions (i.e. lower error rate than required) do not contribute to a higher RMSE.

**Table S1:** Summary of some the features and available configurations within CPSign. Bold faced words are the default for the given row/item in the table. Further note that all of these, save from the predictor type, can be extended and injected with custom implementations. Further note that the item "Data transformations" list types of transformations, and there can be several implementations to chose from from each type.

| **Item** | **Options** | | | | |
|---|---|---|---|---|---|
| Predictor type | **ICP/ACP** | TCP | Venn-ABERS | | |
| Classification scorer models | **LinearSVC** | C_SVC | NuSVC | LogisticRegression | |
| Regression scorer models | **LinearSVR** | EpsilonSVR | NuSVR | | |
| Classification nonconf metrics | **ND2H**[a] | PD2H[b] | InverseProbability | Probability Margin | |
| Regression Nonconf metrics | **LogNormalized** | Normalized | AbsDiff | SignedNormalized | |
| P-value calculation | Standard | **Smoothed** | Linear interpolation | Spline interpolation | |
| Data splitting | **Random** | RandomStratified | Folded | FoldedStratified | PreDefined |
| Data transformations | Duplicate resolver | Filters | Imputation | Feature selection | Feature scaling |
| Descriptors | **Signatures** | ECFP | UserSupplied | CDK descriptors | |

[a] ND2H: Negative Distance to hyperplane
[b] PD2H: Positive Distance to hyperplane

**Table S2:** Performance ranking of the modeling methods in the comparison. The "Top" column shows the number of datasets that each method produced the most efficient predictions, whereas the "Rank" column displays the sum of ranks across all datasets. The best value in each column is displayed in bold. OF: Observed Fuzziness.

| | Classification | | | | | | | | Regression | | | | | |
|---|---|---|---|---|---|---|---|---|---|---|---|---|---|---|
| | OF | | Average C | | | | | | Prediction interval width | | | | | |
| | | | $\varepsilon = 0.1$ | | $\varepsilon = 0.2$ | | $\varepsilon = 0.3$ | | $\varepsilon = 0.1$ | | $\varepsilon = 0.2$ | | $\varepsilon = 0.3$ | |
| | **Top** | **Rank** | **Top** | **Rank** | **Top** | **Rank** | **Top** | **Rank** | **Top** | **Rank** | **Top** | **Rank** | **Top** | **Rank** |
| CPSign | **5** | 44 | **7** | **39** | **5** | **40** | 4 | 48 | 0 | 56 | 1 | 53 | 3 | 53 |
| CPSign tuned | **5** | **39** | 2 | 46 | 3 | 43 | 4 | **40** | **12** | **29** | **11** | **32** | **7** | **31** |
| FP+RF tuned | 1 | 62 | 2 | 65 | 2 | 66 | 0 | 64 | 2 | 72 | 1 | 73 | 2 | 75 |
| Chemprop | 2 | 44 | 2 | 40 | 3 | 44 | **5** | **40** | 1 | 62 | 3 | 58 | 3 | 56 |
| Chemprop tuned | 3 | 51 | 3 | 50 | 3 | 47 | 3 | 48 | 3 | 51 | 2 | 54 | 3 | 55 |

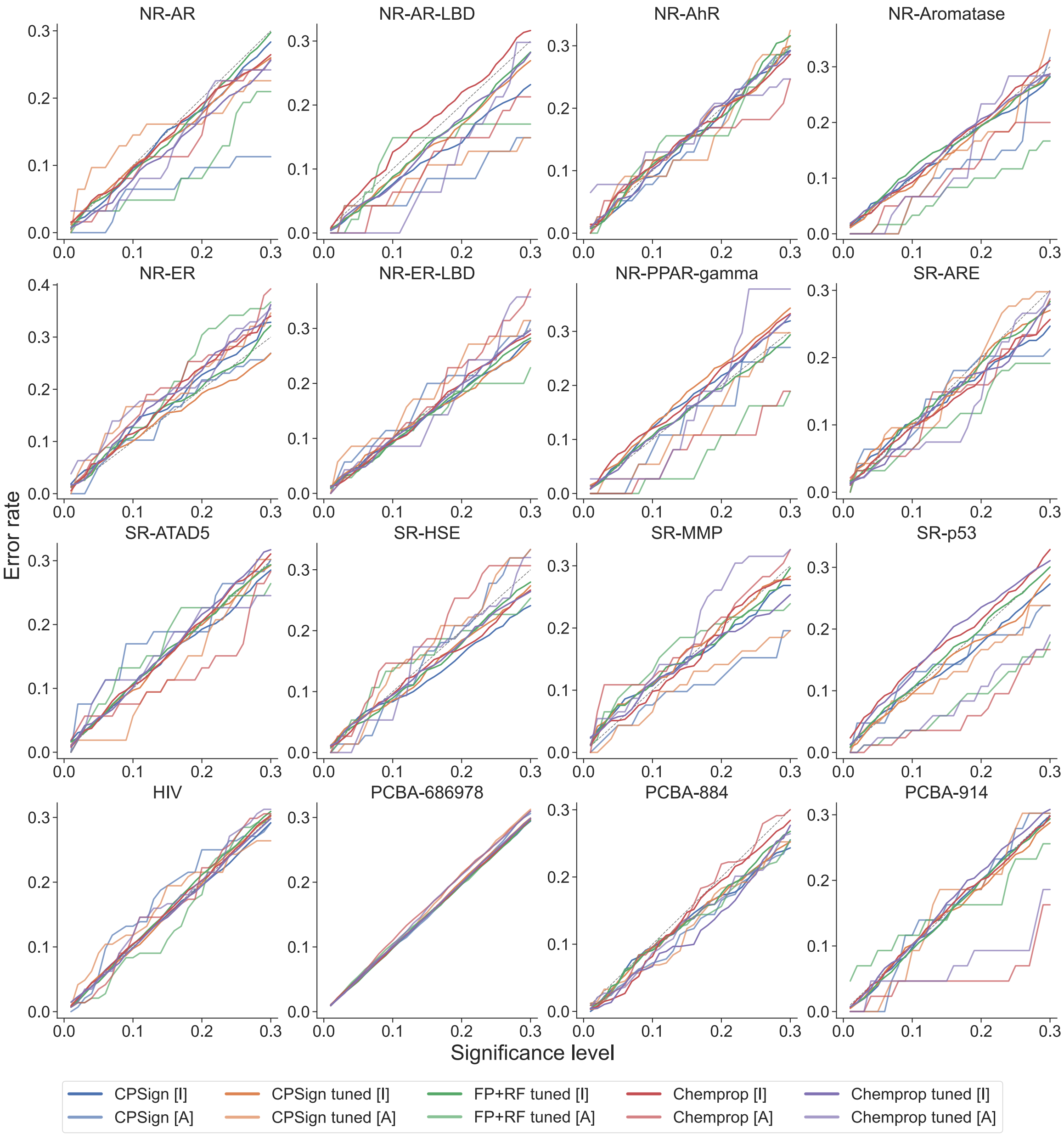


**Figure S2:** Calibration curves for the classification datasets, showing one curve for each class (I: inactive, A: active). The active class is the minority class for all datasets, displaying worse calibration, which can be more easily seen in the aggregation in Figure S1. All calibration curves were evaluated from 0.01,0.02,...,0.3.

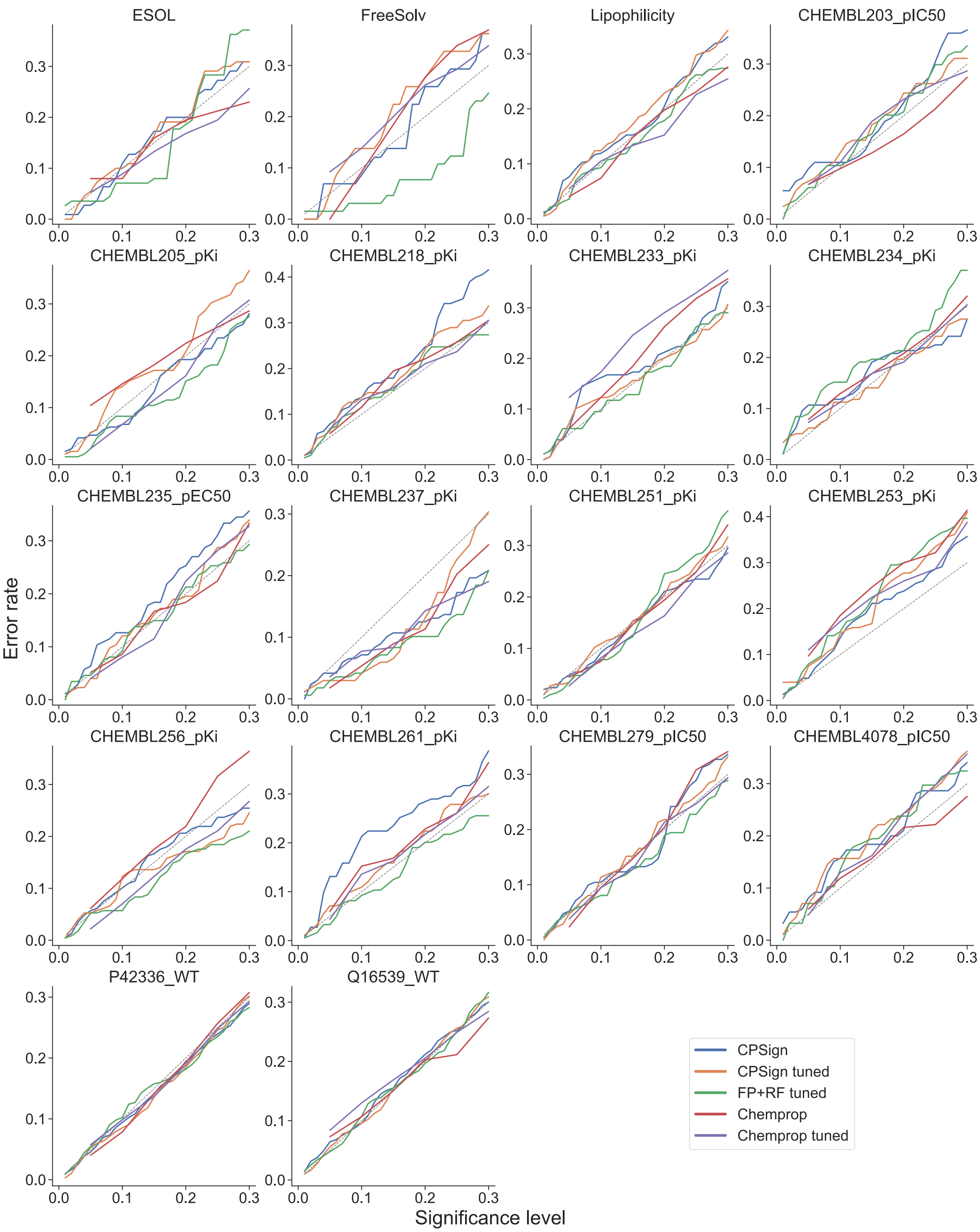


**Figure S3:** Calibration curves for the regression datasets. Note that the Chemprop methods were only evaluated using the six significance levels 0.05,0.1,...,0.3, whereas the other methods used significance levels 0.01,0.02,...,0.3.

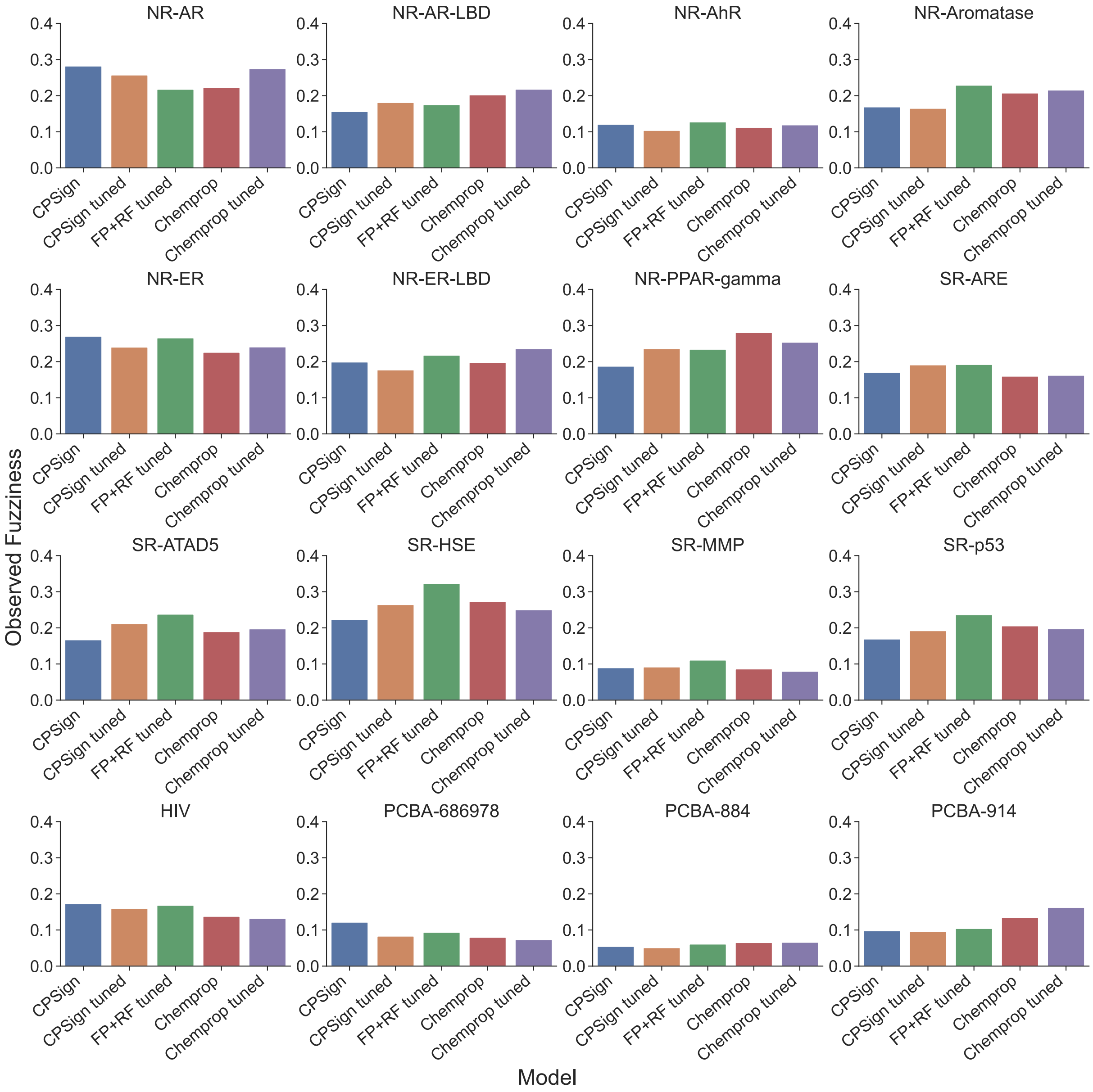


**Figure S4:** Comparison of the Observed Fuzziness (OF) of the evaluated methods for all sixteen datasets. A lower OF score is preferable. Larger differences can be seen when analyzing the datasets individually, and each modeling method produces the best predictions for at least one datase (see Table S2).

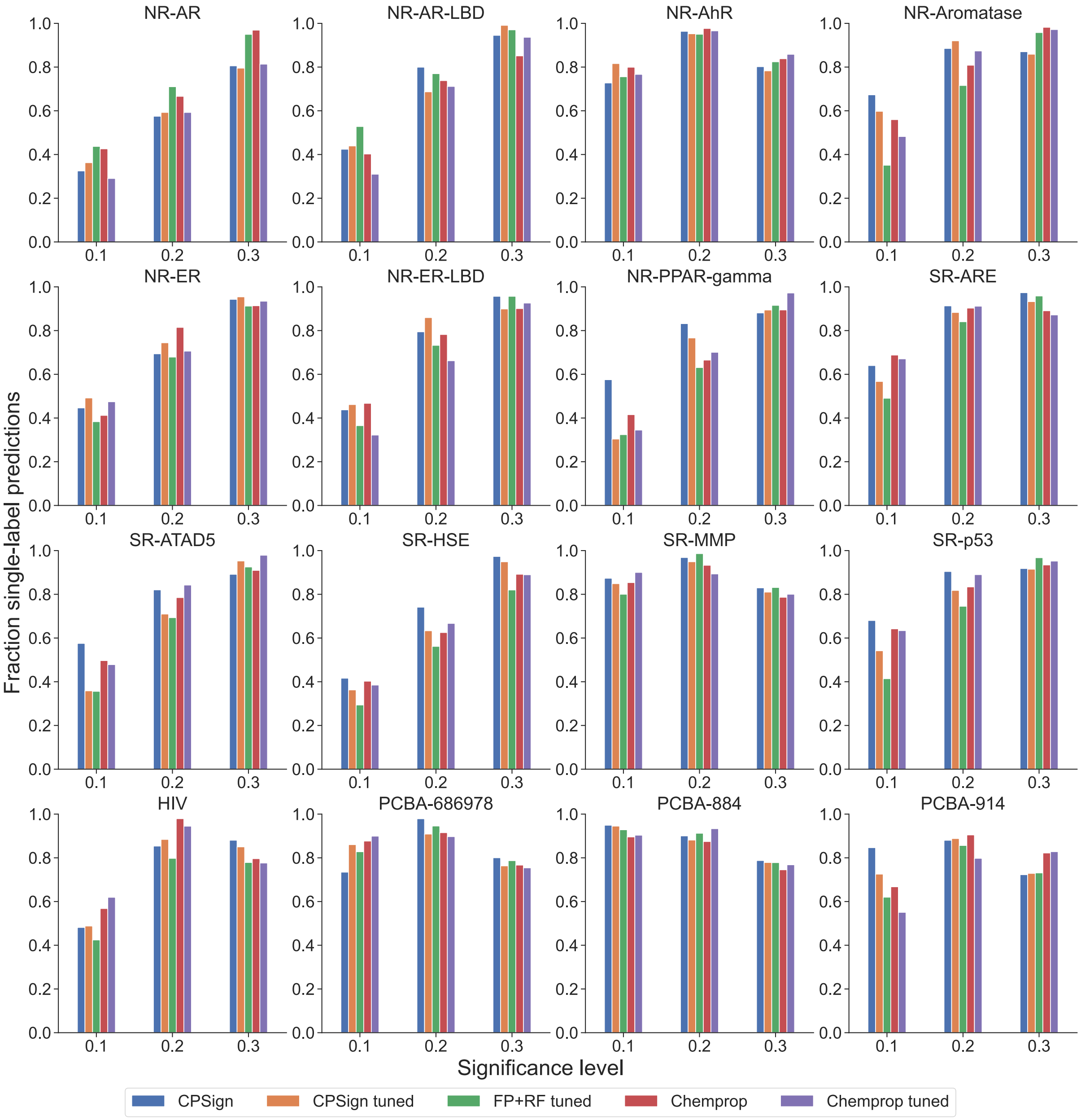


**Figure S5:** Median fraction of single-label predictions for the evaluated methods for all sixteen datasets, based on the three significance levels 0.1, 0.2 and 0.3 (corresponding to confidence levels 90 %, 80 % and 70 %, respectively). A higher fraction is preferable. Similarly as Figure S4 the prediction results display larger differences than the aggregated results, with different methods being the top-performing one.

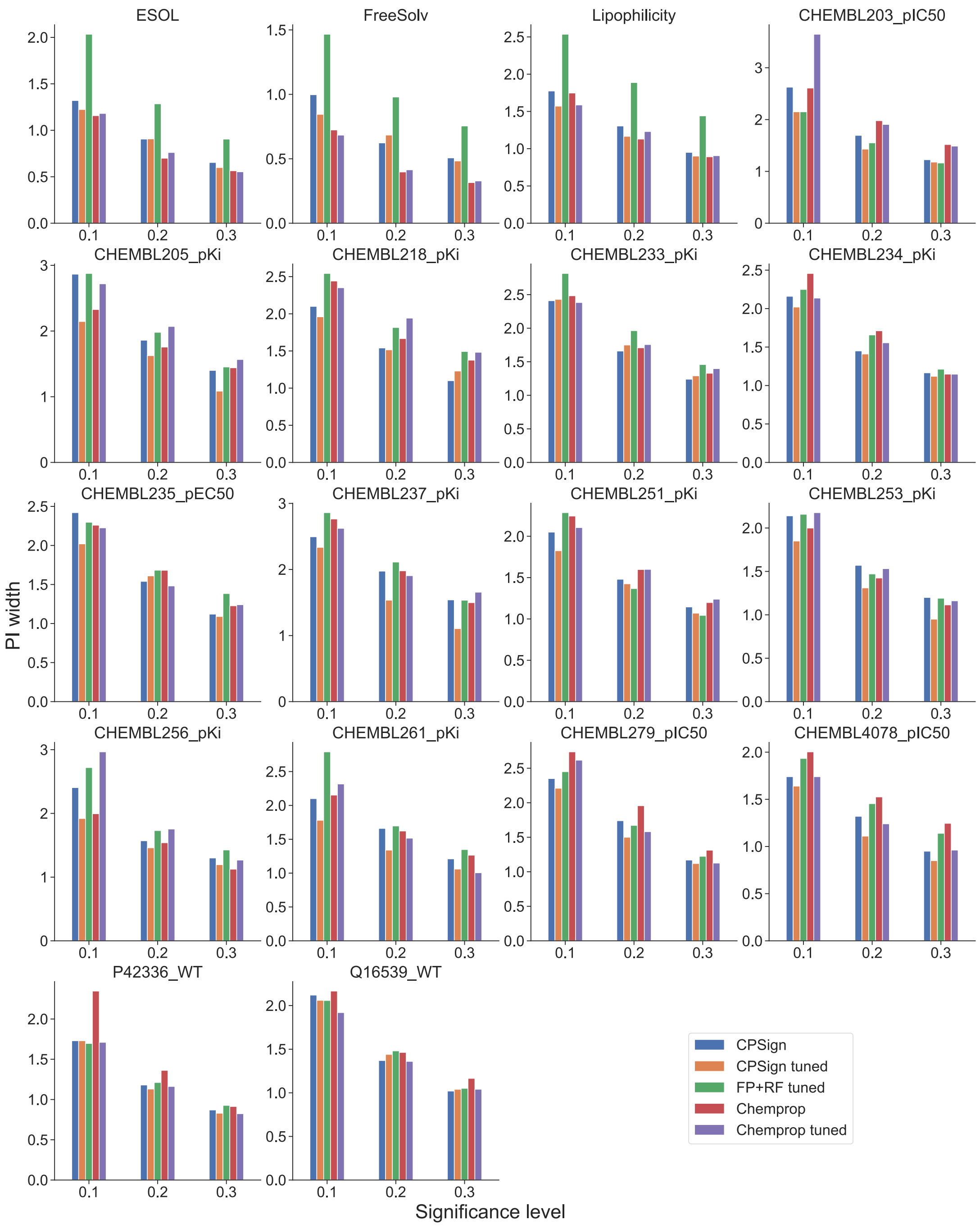


**Figure S6:** Median prediction interval (PI) width for all regression datasets, for significance levels 0.1, 0.2 and 0.3 (corresponding to confidence levels 90 %, 80 % and 70 %, respectively). A lower value is preferable (i.e. tighter prediction intervals).

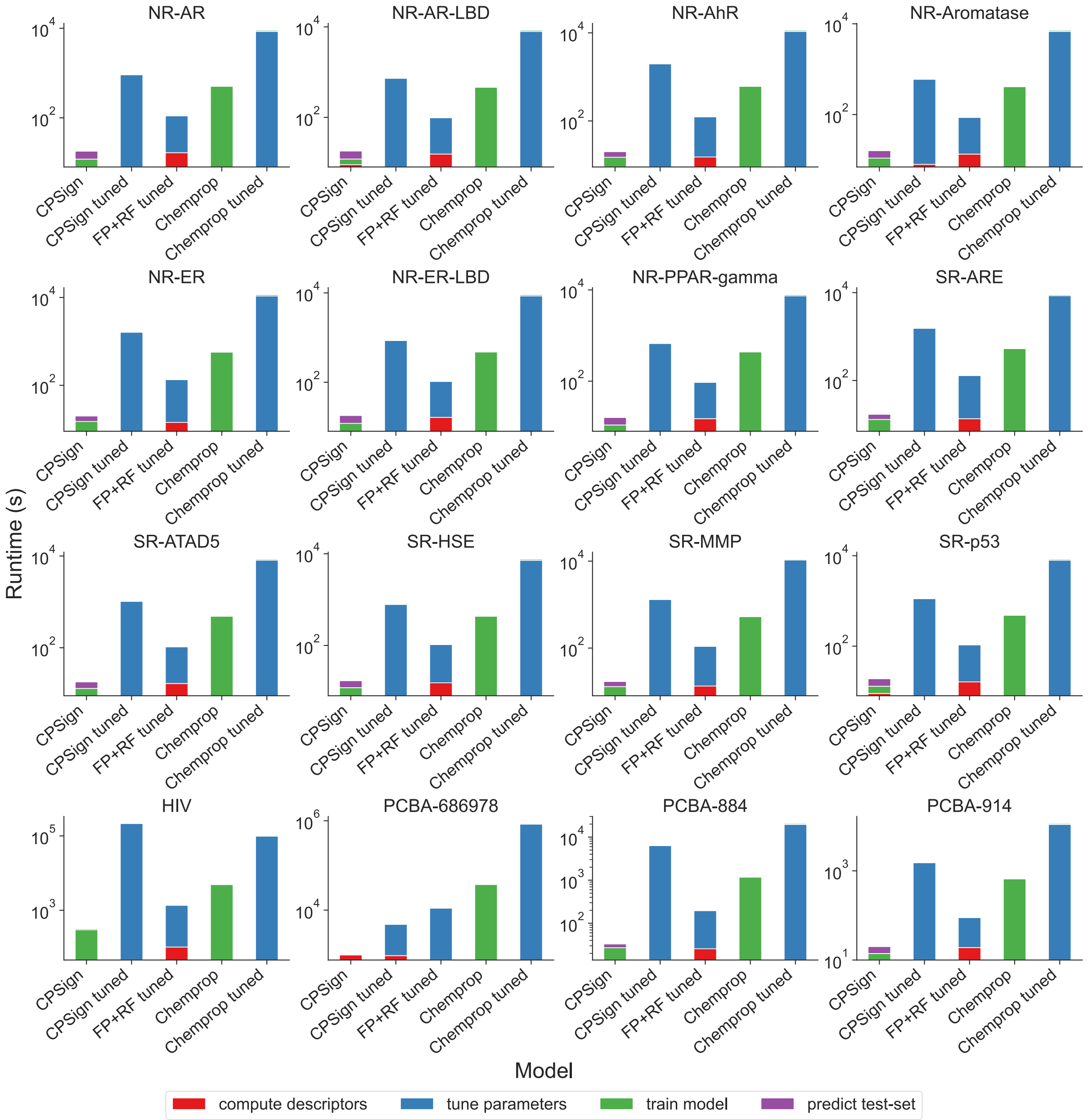


**Figure S7:** Runtime comparison for individual classification datasets, using a logarithmic y-axis. The relative runtimes are consistent across all runs, except the PCBA-686978 dataset (where CPSign and CPSign tuned were run with linear SVM kernels). Note that the two Chemprop methods do not contain a separate step for computing descriptors, which instead is included in the tuning and training steps.

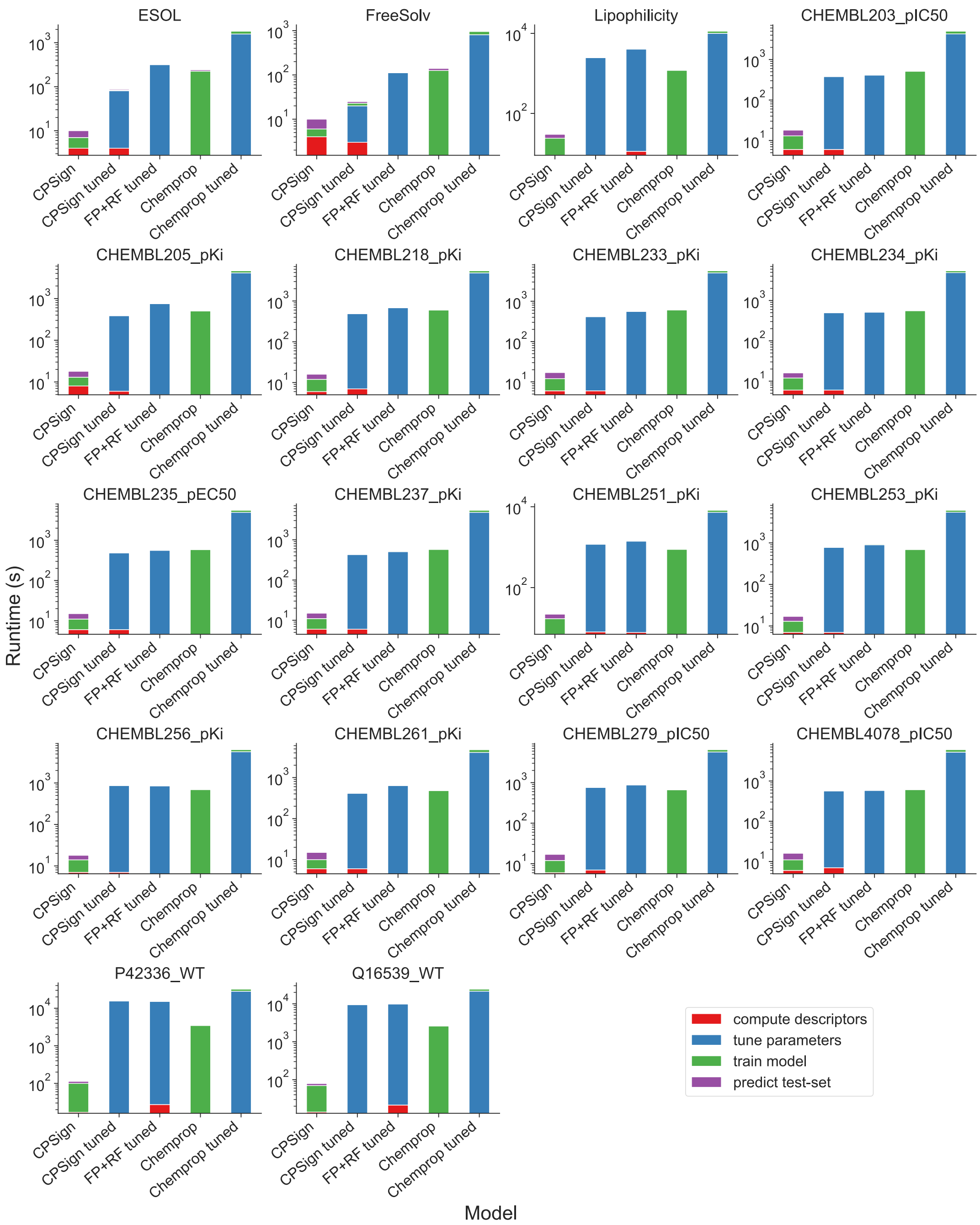


**Figure S8:** Runtime comparison for individual regression datasets, using a logarithmic y-axis. The relative runtimes are consistent across all runs, with only minor differences between the CPSign tuned, FP+RF tuned and Chemprop methods. Note that the two Chemprop methods do not contain a separate step for computing descriptors, which instead is included in the tuning and training steps.
